## Supplementary File 1 for "The substrate-binding domains of the osmoregulatory ABC importer OpuA transiently interact"

### Supplementary Note 1: optimization of purification and nanodisc reconstitution

In order to obtain a homogenous nanodisc sample, which consists of a tetrameric OpuA complex in a lipid-filled nanodisc, the previous protocol was optimized on several aspects; the detailed procedure is describe in Supplementary Note 2. Once OpuA is solubilized with 0.5% DDM (w/w) in the presence of 20% glycerol (v/v) it is necessary to prevent dissociation of the complex [1]. Therefore, both the vesicle solubilization step and the incubation of the sample with the Ni-Sepharose resin were shortened to 30 min. Based on the shape of the size-exclusion chromatography (SEC) profile this new procedure reduced loss of OpuAA subunits **(Figure S1)**.

Knowing that the OpuA complex is not very stable and that high pressure lysing of cells exerts stress on proteins as well, 20% glycerol (v/v) was added during the preparation of membrane vesicles. Again, on the basis of the SEC profile this markedly reduced the loss of OpuAA subunits **(Figure S2)**.

Shaking or agitation imposes potentially harmful air-water interface forces on proteins. Therefore, agitation during membrane vesicle solubilization and Ni-Sepharose binding was prevented in the new method. Also, less imidazole was used for the elution of the OpuA complex. Furthermore, we removed the His-tag from the MSP proteins, utilizing the TEV-cleavage site that is located between MSP-protein and the His-tag. We removed empty nanodiscs by an extra IMAC step after overnight reconstitution.

For the preparation of nanodiscs we switched from MSP1D1 to MSP1E3D1, because cryoEM structures indicate that there is limited space for lipids around OpuA in MSP1D1 nanodiscs [2]. Previously, the synthetic lipid mixture was solubilized prior to reconstitution by extrusion through a 400 nm filter and subsequent vortexing in the presence of 12 mM DDM. The lipids were diluted to the proper concentration in the reconstitution mixture, but the DDM concentration was kept at 12 mM. A high DDM concentration during reconstitution induce dissociation of OpuA complexes **(Figure S3,S4)**. Therefore, in the new method no extra DDM was added when the lipids were diluted in the reconstitution mixture. Consequently, the DDM concentration in the reconstitution mixture was less than 8 mM.

During previous reconstitutions, the molar ratio of OpuA:MSP:lipid was 1:20:1000. However, after extrusion through a 400 nm filter and vortexing of the synthetic lipids in the presence of 12 mM DDM, the solution still appeared turbid. This could be an indication for the presence of unsolubilized liposomes. Those lipids will not participate in the reconstitution process and can contribute to heterogeneity in the final nanodisc sample. In the current method, the lipids are sonicated in 50 mM KPi pH 7.0 at a concentration of 6.25 mg/mL. Afterwards, the sample is incubated for at least 2 hours with 1% DDM (w/w). The lipid sample becomes fully transparent before it is used for the nanodisc reconstitution of OpuA.

Because most, if not all, lipids are solubilized with this new method and another membrane scaffolding protein (MSP1E3D1) is used, the molar ratio of OpuA:MSP:lipid was re-optimized. Since a OpuA:MSP ratio of 1:20 created a lot of empty nanodiscs, the ratio was changed to 1:10. The OpuA:MSP:lipid ratio of 1:10:200 gave the best SEC profile in terms of monodispersity and width of the SEC profile **(Figure S5).** For more details on the new purification procedure, see Supplementary note 2. For a short overview of the optimization steps, see **Table S1**.

### Supplementary Note 2: detailed purification procedure

**Preparation of crude membrane vesicles • TIMING 7-9 h**

1. Use cells with an $OD_{600}$ of 150-200, which are resuspended in 50 mM KPi pH 7.5 plus 20% (v/v) glycerol. Perform all steps of this procedure on ice or in a cooled environment (4 to 8 °C), unless otherwise indicated.

▲CRITICAL STEP The presence of glycerol throughout the preparation of crude membrane vesicles is necessary to prevent dissociation of OpuAA subunits from the full OpuA complex.

2. Add 100 µg/ml DNAse plus 2 mM $MgSO_4$. Also slowly add 1 mM PMSF, dissolved in isopropanol, while stirring the cell suspension.
3. Rupture the cells by passing them through a cooled (4 °C) high-pressure lyser HPL6 (Maximator GmbH) at a pressure of 30 kPsi. Repeat this step a second time.
4. Add 5 mM of EDTA and centrifuge at 4 °C for 25 min, 15,000 xg in Beckman J-20XP, using the JA-25.50 rotor.
5. Transfer the supernatant to Type 45 Ti or Type 50.2 Ti Bottle Assembly tubes by pouring.

▲CRITICAL STEP The pellet can easily come loose. When pellet material comes along during the transfer of the supernatant, repeat the previous centrifuge step.

6. Centrifuge at 4 °C for 135 min, 42,000 rpm in Beckman Optima XE-90, using the Type 45 Ti rotor (138,298 xg at $r_{av}$); or at 4 °C for 75 min, 45,000 rpm in Beckman Optima XE-90, using the Type 50.2 Ti rotor (184,842 xg at $r_{av}$).
7. Remove the supernatant by pouring and release the pellet from the tube by using a pestle from a potter tube and a few mL of 50 mM KPi pH 7.5 plus 20% glycerol (v/v).
8. Pour the pellet material in a potter tube and wash the centrifuge tube at least two times with 50 mM KPi pH 7.5 plus 20% glycerol (v/v) to retrieve all the material.
9. Fill the potter tube with extra buffer to obtain a volume that is equal to approximately 50% of the lysate after disruption, and resuspend the membranes completely by using the pestle of the potter tube.
10. Repeat step 5-8, but now use only 6 mL of 50 mM KPi (pH 7.5) plus 20% (v/v) glycerol per 100 mL of cells with an $OD_{600}$ of 150-200 (volume of the cells at the start of the procedure) to resuspend the membrane vesicles.
11. Determine the protein concentration by means of a Pierce BCA assay. The expected concentration is 8-16 mg/mL.
12. Divide the sample in aliquots of 18 mg membrane vesicles, flash freeze the aliquots in liquid nitrogen and store them at -80 °C.

**Solubilization of synthetic lipids • TIMING 30 min**

13. Thaw 250 µL of 25 mg/mL lipids in 50 mM KPi pH 7.0. Dilute the lipids, using the same buffer, to 6.25 mg/mL in a 15 mL Falcon tube at room temperature.
14. Tip-sonicate the lipids in an ice-water bath for 8 cycles of 15 sec with a 45 sec interval at an amplitude of 77 µm.

▲CRITICAL STEP For this synthetic lipid mix, tip-sonication is necessary to create small unilamellar vesicles, because DDM is not able to fully solubilize multilamellar vesicles within a time period of less than 5 h.

15. Transfer 900 µL of the sonicated lipids to a small glass vial, and incubate at room temperature with 100 µL of 10% DDM (w/w) until the lipids are needed for OpuA reconstitution in lipid nanodiscs.

**Solubilization of crude membrane vesicles • TIMING 1 h**

16. Thaw the crude membrane vesicles at room temperature but put them on ice as soon as they are thawed.
17. Dilute the crude membrane vesicles on ice in MLA-80 centrifuge tubes to 3 mg/mL in 6 mL. The final mixture should contain 50 mM KPi pH 7.0-7.5, 20% (v/v) glycerol, 200 mM KCl plus 0.5% (w/w) DDM. As an example, 9 mg/mL membrane vesicles are diluted according to the pipetting scheme below:

| Reagent | Stock concentration | Volume (µL) |
|---|---|---|
| MQ | - | 700 |
| Glycerol | 50% (v/v) | 1600 |
| KPi pH 7.0 | 200 mM | 1000 |
| KCl | 3 M | 400 |
| DDM | 10% (w/w) | 300 |
| Membrane vesicles* | 9 mg/mL | 2000 |

* Take into account that membrane vesicles are already in a 50 mM KPi plus 20% (v/v) glycerol.

18. Incubate the mixture for 30 min on ice and mix once by pipetting after 15 min.

▲CRITICAL STEP The sample is not mixed by agitation, because this induces dissociation of OpuAA subunits from the full OpuA complex.

19. Centrifuge at 4 °C for 20 min, 80,000 rpm in Beckman MAX-E, using the MLA-80 rotor (336,896 xg at $r_{av}$).

**Purification ($Ni^{2+}$-Sepharose affinity chromatography) • TIMING 2 h**

20. Mix the supernatant in a 10 mL disposable chromatography column with 10 mM imidazole pH 7.5, and 0.5 mL of $Ni^{2+}$-Sepharose (bed volume) that was washed with 12 column volumes of MQ and equilibrated with 4 column volumes of 50 mM KPi pH 7.0, 200 mM KCl, 20% (v/v) glycerol, 0.04% (w/w) DDM (Buffer A) plus 10 mM imidazole. Mix by gentle pipetting with a 1 mL pipette and make sure that the outlet of the column is closed.
21. Allow the $Ni^{2+}$-Sepharose to sediment in 10-20 min, before mixing the sample a second time.

▲CRITICAL STEP The sample is not mixed by agitation, because this induces dissociation of OpuAA subunits from the full OpuA complex.

22. Drain the column once the $Ni^{2+}$-Sepharose is sedimented again and wash twice with 10 column volumes of buffer A plus 50 mM imidazole pH 7.5.
23. Elute OpuA by adding consecutively one time 0.6 and four times 0.4 column volumes of buffer A plus 200 mM imidazole pH 7.5. Collect the five fractions separately. Most protein is expected in the third and fourth fraction.

**Nanodisc reconstitution and second purification • TIMING 16-18 h**

24. Mix the following components in a 1.5 mL Eppendorf tube to an end volume of 700-900 µL and agitate for 1 hour at 4 °C:

| Reagent | Stock concentration | End concentration |
|---|---|---|
| MQ | - | - |
| KPi pH 7.0* | 1 M | 50 mM |
| Lipids | 5.63 mg/mL | 900 µM |
| MSP1E3D1 | 140-160 µM | 45 µM |
| OpuA | 13-23 µM | 4.5 µM |

* Take into account that OpuA and the lipids are already in 50 mM KPi pH 7.0

▲CRITICAL STEP This agitation step is necessary for nanodisc reconstitution but not ideal for OpuA. Therefore, try to minimize the empty volume in the tube.

25. Add 500 mg of semi-dry activated SM-2 Bio-Beads to the sample and incubate overnight under gentle agitation at 4 °C.
26. Separate the nanodiscs from the beads by using a syringe with a needle (⌀ 0.2 mm). To prevent transfer of beads that are stuck inside the needle, remove the needle before transferring the sample from the syringe in a new 1.5 mL Eppendorf tube.
27. Spin down large particles at 20,817 xg for 15 min at 4°C.

▲CRITICAL STEP It is possible to directly proceed to size-exclusion chromatography if the resolution of the column is good enough to separate empty nanodiscs from the nanodiscs that contain the protein of interest. In this study, point 28-31 are mainly used for maleimide based cysteine labeling.

28. Pipette the supernatant gently to a 2 mL disposable chromatography column, which contains 0.2-0.3 mL of $Ni^{2+}$-Sepharose (bed volume) that was washed with 10 column volumes of MQ and equilibrated with 5 column volumes of 50 mM KPi pH 7.0.
29. Let the sample flow through, collect the flow-through as a single fraction and reapply it twice back to the $Ni^{2+}$-Sepharose.
30. Wash two times with 5 column volumes of 20 mM HEPES-K pH 7.0, 300 mM KCl plus 25 mM imidazole pH 7.5.
31. Elute OpuA by adding consecutively 0.8 and 2.5 column volumes of 20 mM HEPES-K pH 7.0, 300 mM KCl plus 200 mM imidazole pH 7.5. Collect the two fractions separately.

**Size-exclusion chromatography and storage • TIMING 1 h**

32. Load the second elution fraction on a Superdex™ 200, 10/300GL Increase column, preequilibrated with 20 mM HEPES-K pH 7.0 plus 300 mM KCl, and collect the elution in 0.5 mL fractions.
33. Take the two best fractions, flash freeze them in liquid nitrogen in 50-100 µL aliquots and store them at -80 °C.

Table S1. Changes in purification and reconstitution conditions, and their effects on the stability or activity of OpuA

| **Old condition** | **New condition** | **Difference (new *versus* old)** |
|---|---|---|
| Prepare crude membrane vesicles in the absence of glycerol | Prepare crude membrane vesicles in the presence of 20% glycerol (v/v) | Less dissociated complex |
| Membrane vesicle solubilization for 1 hour | Membrane vesicle solubilization for 30 min | Less dissociated complex |
| Agitation during vesicle solubilization and binding of OpuA to $Ni^{2+}$-Sepharose | Incubation without agitation | Not tested |
| Use 500 mM imidazole for elution | Use 200 mM imidazole for elution | Not tested |
| MSP with His-tag | MSP without His-tag | None |
| Use MSP1D1 | Use MSP1E3D1 | None |
| No IMAC after reconstitution | IMAC after reconstitution | Removal of empty nanodiscs and cleaner SEC profile |
| Add to 12 mM DDM during reconstitution | Add ≤ 8 mM DDM during reconstitution | Less dissociated complex |
| Use extruded lipid mixture for reconstitution | Use sonicated lipid mixture for reconstitution | More reproducible SEC profiles. Smaller shoulder on the left in the SEC profile |
| OpuA:MSP:lipid is 1:20:1000 (molar ratio) during nanodisc reconstitution | OpuA:MSP:lipid is 1:10:200 (molar ratio) during nanodisc reconstitution | Reproducible and monodispersed SEC profile |

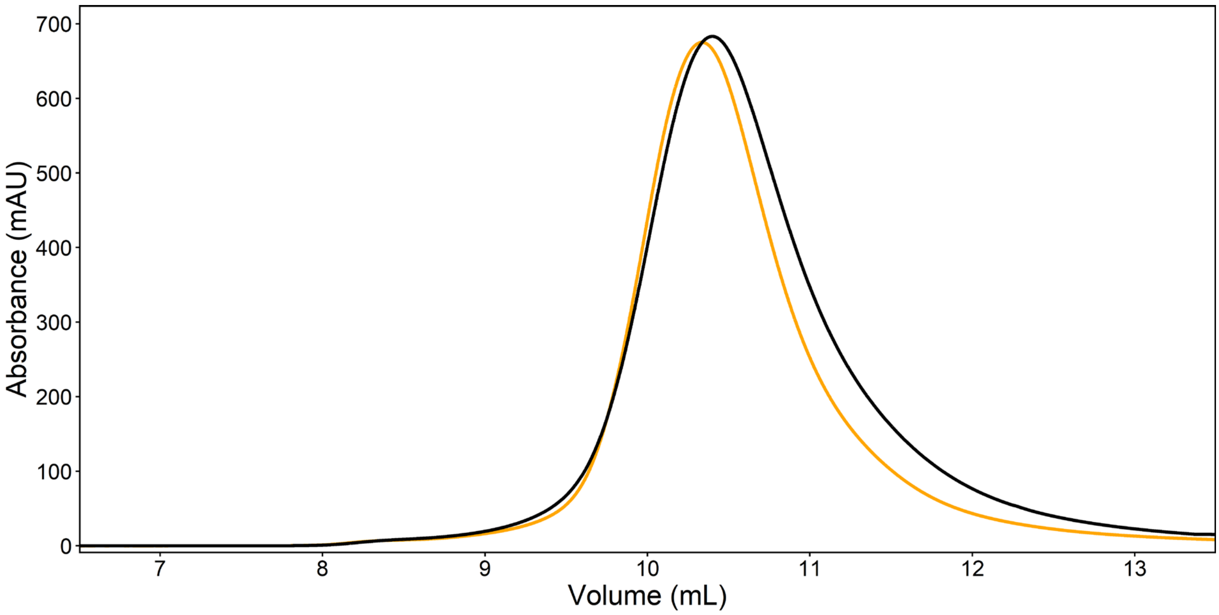


FIGURE S1. Size-exclusion profiles of purified OpuA in DDM. Previously, the membrane vesicle bearing OpuA were solubilized for one hour in 50 mM KPi pH 7.0, 200 mM KCl, 0.5% DDM (w/w) plus 20% glycerol (v/v) and subsequently incubated for 1 hour with Ni-Sepharose resin (black line). Both steps were shortened to 30 min (yellow line).

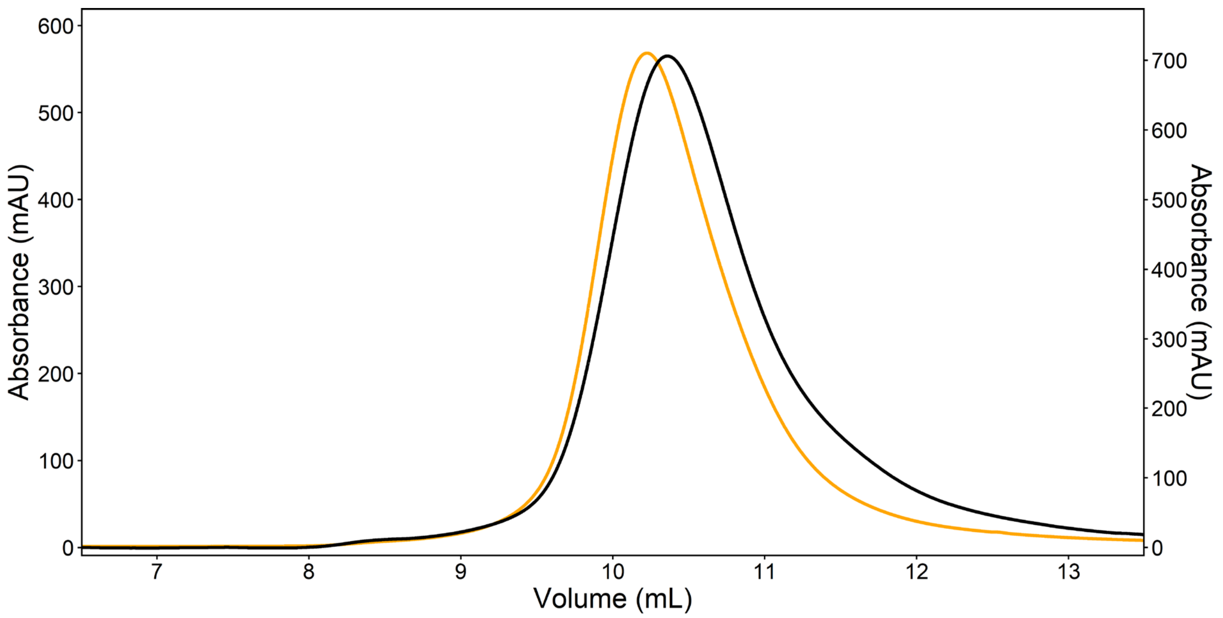


FIGURE S2. Size-exclusion profiles of purified OpuA in DDM. Previously, *L. lactis* cells were lysed in 50 mM KPi pH 7.5 without glycerol (black line, right y-axis). In the new method, 20% glycerol (v/v) was added during high-pressure lysis (yellow line, left y-axis).

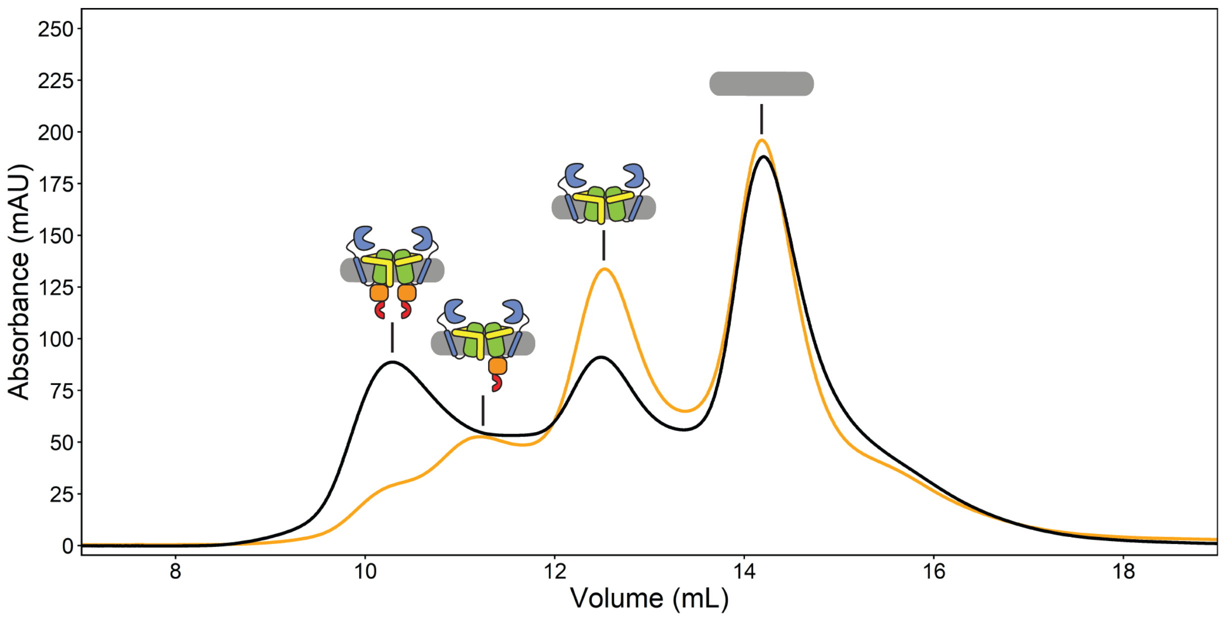


FIGURE S3. Size-exclusion profiles of purified OpuA in MSP1D1 nanodiscs. Before the addition of Bio-Beads, the reconstitution mixture contained either 12 (black line) or 39 (yellow line) mM DDM.

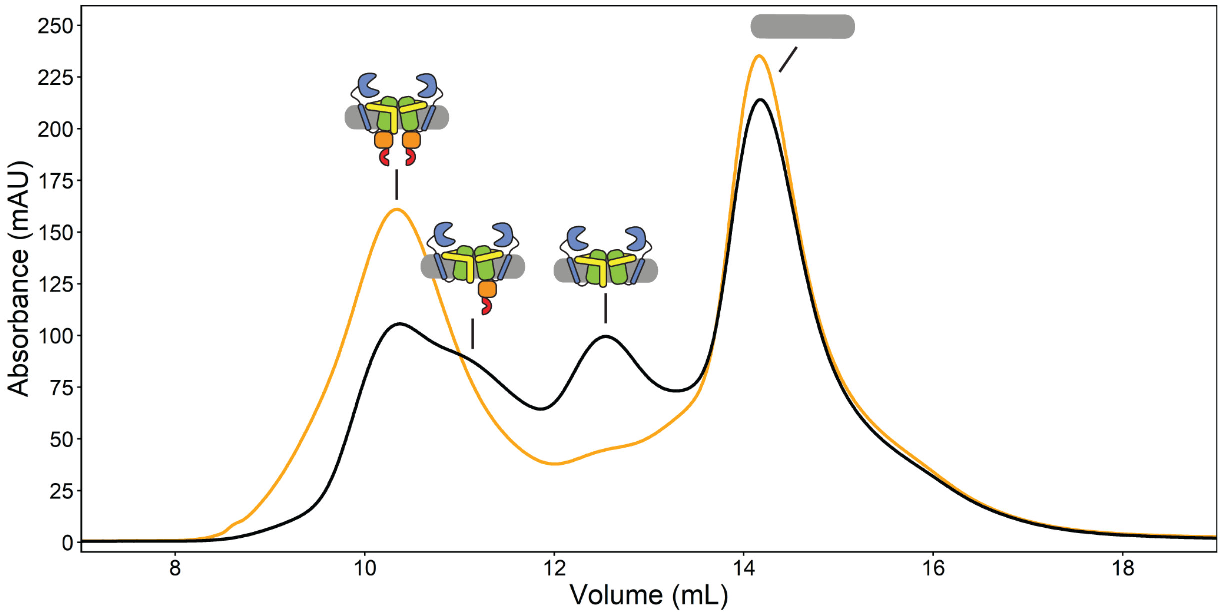


FIGURE S4. Size-exclusion profiles of purified OpuA in MSP1D1 nanodiscs. Before the addition of Bio-Beads, the reconstitution mixture contained either 12 (black line) or 8 (yellow line) mM DDM.

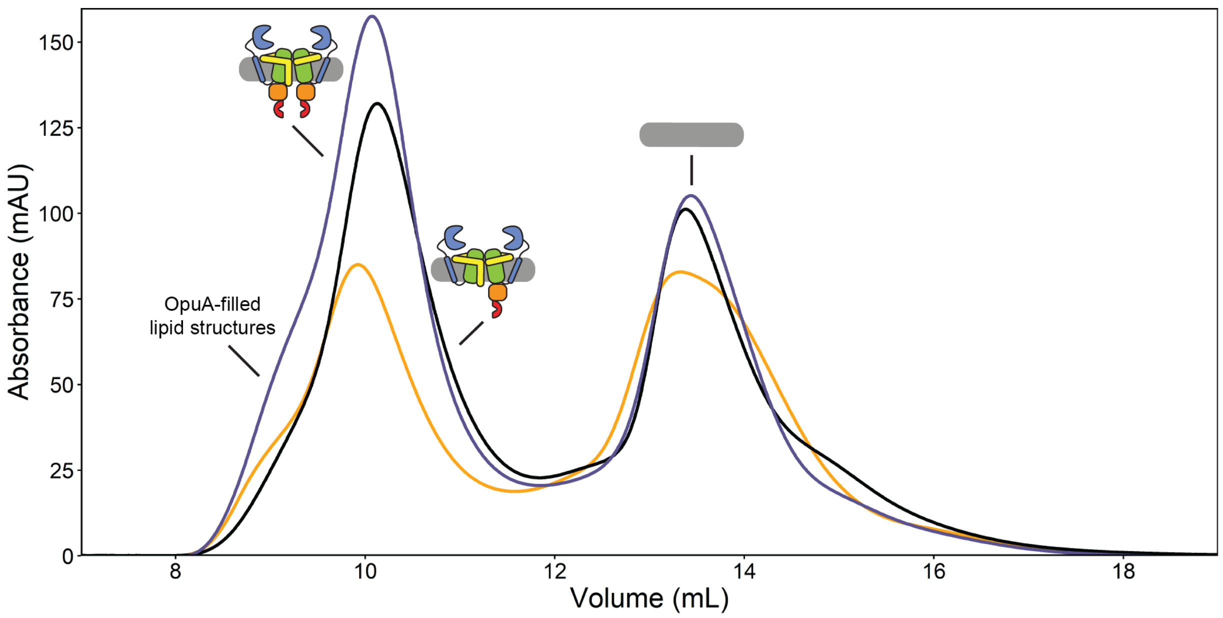


FIGURE S5. Size-exclusion profiles of purified OpuA in MSP1E3D1 nanodiscs with different OpuA:MSP1E3D1:lipid molar ratios. The OpuA:MSP1E3D1:lipid ratios are 1:10:200 (black), 1:10:300 (purple) and 1:10:400 (yellow).
