## Supplementary File 2 for "The substrate-binding domains of the osmoregulatory ABC importer OpuA transiently interact"

Table S1. ATPase activity of OpuA variants K521C and N414C before and after labeling with maleimide dyes†

| | **Activity unlabeled variant ($min^{-1}$)** | **Activity labeled variant ($min^{-1}$)** |
|---|---|---|
| K521C | 342 +/- 71ǂ | 263 +/- 52 |
| N414C | 332 +/- 25 | 246 +/- 55 |

† Buffer conditions: 50 mM HEPES-K pH 7.0, 100 µM glycine betaine, 10 mM Mg-ATP plus 600 mM KCl

ǂ The errors refer to the standard deviation over at least two measurements with different protein purifications and membrane reconstitutions, each consisting of three technical replicates.

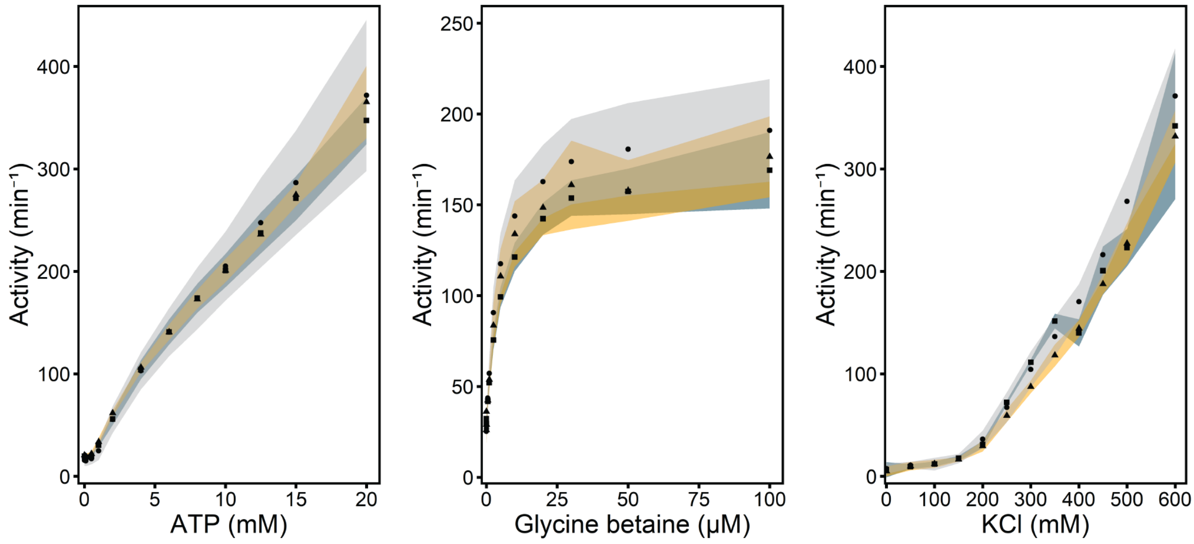


FIGURE S1. Enzyme-coupled ATPase assay of wildtype OpuA (circles, grey shading), OpuA-K521C (squares, blue shading) and OpuA-N414C (triangles, yellow shading). A standard sample contains 50 mM HEPES-K pH 7.0, 450 mM KCl, 20 mM Mg-ATP, 100 µM glycine betaine, 4 mM phospho*enol*pyruvate, 600 µM NADH, 2.1 to 3.5 U of pyruvate kinase plus 3.2 to 4.9 U of lactate dehydrogenase. Standard deviation over at least two measurements with different protein purifications and membrane reconstitutions, each consisting of three technical replicates is represented as shaded areas.

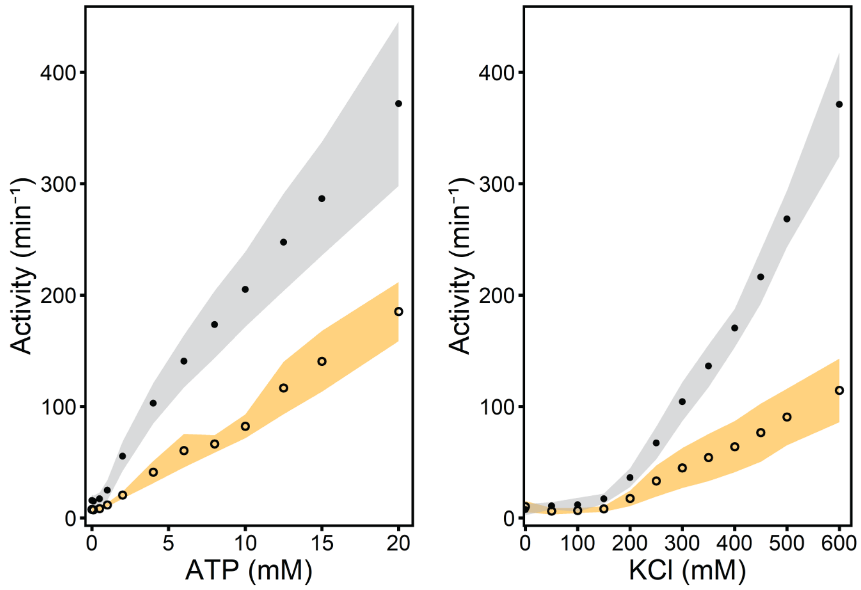


FIGURE S2. Enzyme-coupled ATPase assay for wildtype OpuA (squares, grey shading) and OpuA-V149Q-K521C (circles, yellow shading). A standard sample contains 50 mM HEPES-K pH 7.0, 450 mM KCl, 20 mM Mg-ATP, 100 µM glycine betaine, 4 mM phospho*enol*pyruvate, 600 µM NADH, 2.1 to 3.5 U of pyruvate kinase plus 3.2 to 4.9 U of lactate dehydrogenase. Standard deviation over at least two measurements with different protein purifications and membrane reconstitutions, each consisting of three technical replicates is represented as
