## Supplementary File 3 for "The substrate-binding domains of the osmoregulatory ABC importer OpuA transiently interact"

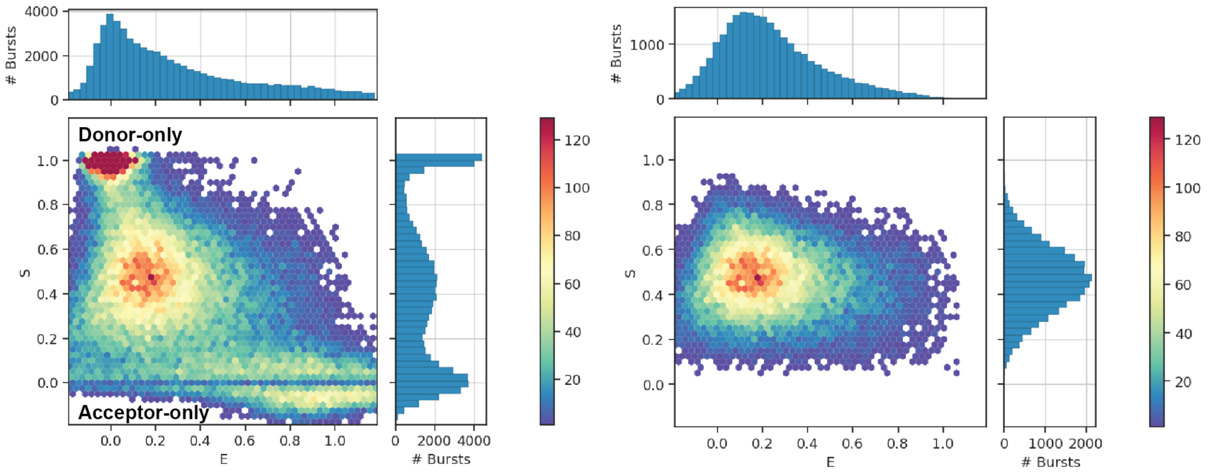


FIGURE S1. Typical 2D E-S histograms of FRET bursts after γ-correction and correction for leakage and crosstalk. Graphs represent data of OpuA-K521C in 50 mM HEPES-K pH 7.0, 600 mM KCl. Left graph shows the bursts after burst selection with a cut-off of minimally 35 photons per burst. Right graph shows the bursts after removal of donor-only and acceptor-only bursts. To filter out acceptor-only bursts, a threshold of minimally 15 photons after donor excitation was set. To filter out donor-only bursts, a threshold of minimally 15 photons was set. Similar graphs for all other OpuA variants and conditions can be found in the publicly available python notebooks (https://doi.org/10.34894/GSIEBW).
