## Supplementary File 4 for "The substrate-binding domains of the osmoregulatory ABC importer OpuA transiently interact"

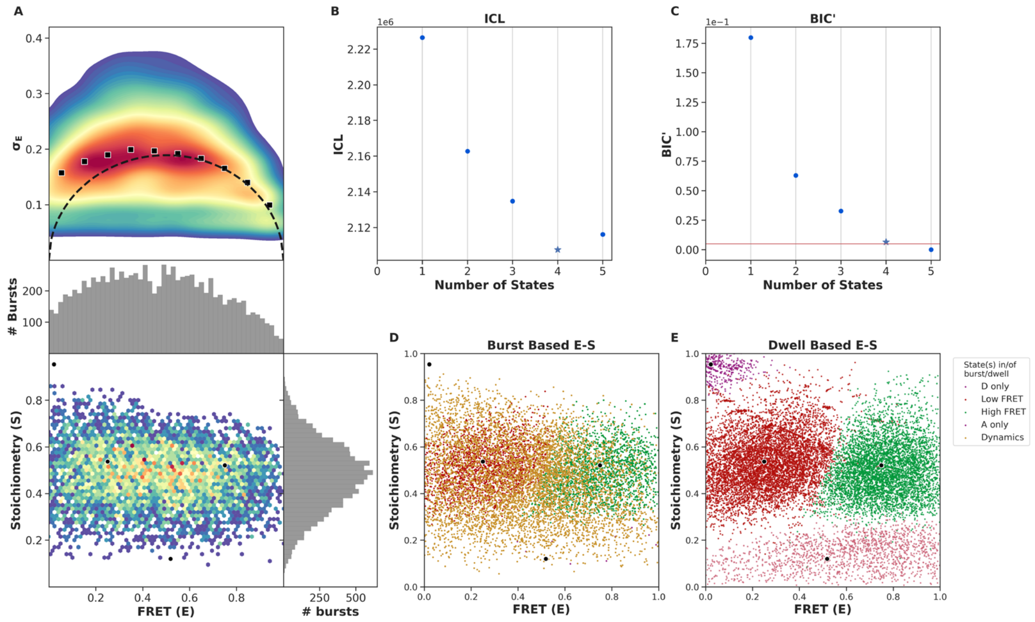


FIGURE S1. Burst variance analysis and mpH2MM of the FRET bursts of OpuA-K521C in 50 mM BIS-TRIS pH 7.0, 0 mM KCl. (A) From top to bottom: (1) Burst variance analysis of the bursts which were corrected by the leakage, crosstalk, and γ-correction factors and which were selected after removing donor-only and acceptor-only bursts. The standard deviation of FRET in each burst is plotted against its mean FRET. Black squares show average values per FRET bin. Black dotted line shows the expected standard deviation in the absence of within-burst dynamics. (2) 2D E-S histogram shows the same data as in (1), with on both sides a histogram that represents the same bursts. (B) Plot of the ICL-values for each final model. The model used in the downstream analysis and following figures is shown as a star. (C) Plot of the BIC'-values for each final model. The red line represents a 0.05 cut-off. The model used is shown as a star. (D) Burst-based 2D E-S scatter plot. Bursts are colored on the basis of the assigned state of the chosen mpH2MM model. If a burst contains more than one state, it is assigned as being dynamic. (E) Dwell-based 2D E-S scatter plot. Dwells are colored on the basis of the assigned state of the chosen mpH2MM model. The dwells were corrected for leakage, direct excitation and the γ-factor. Black dots in A, D and E represent the average value of each state.

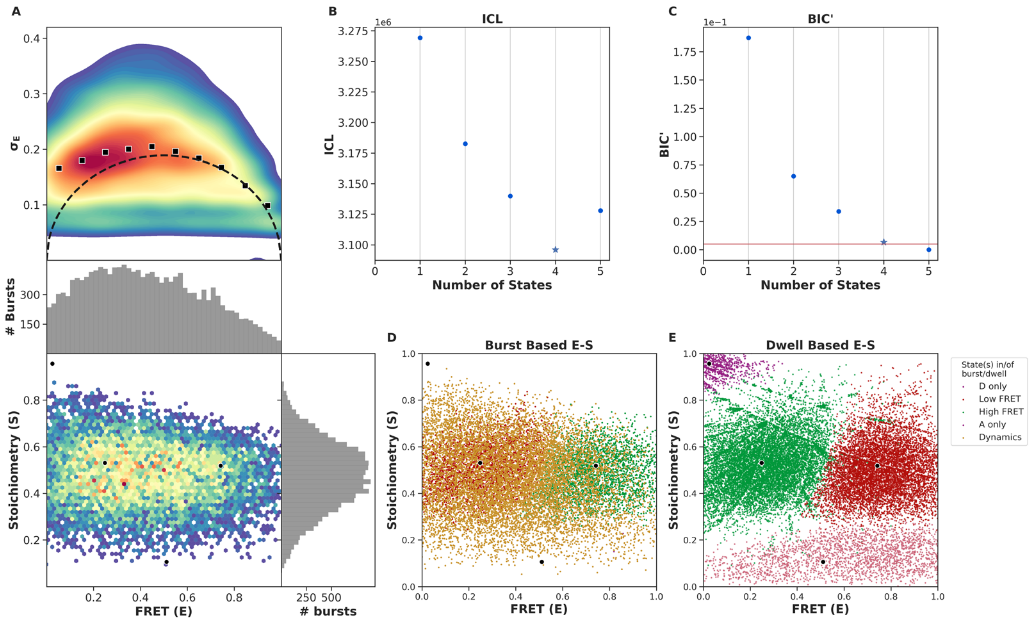


FIGURE S2. Burst variance analysis (BVA) and mpH2MM of the FRET bursts of OpuA-K521C in 50 mM HEPES-K pH 7.0, 0 mM KCl. (A) From top to bottom: (1) Burst variance analysis of the bursts which were corrected by the leakage, crosstalk, and γ-correction factors and which were selected after removing donor-only and acceptor-only bursts. The standard deviation of FRET in each burst is plotted against its mean FRET. Black squares show average values per FRET bin. Black dotted line shows the expected standard deviation in the absence of within-burst dynamics. (2) 2D E-S histogram shows the same data as in (1), with on both sides a histogram that represents the same bursts. (B) Plot of the ICL-values for each final model. The model used in the downstream analysis and following figures is shown as a star. (C) Plot of the BIC'-values for each final model. The red line represents a 0.05 cut-off. The model used is shown as a star. (D) Burst-based 2D E-S scatter plot. Bursts are colored on the basis of the assigned state of the chosen mpH2MM model. If a burst contains more than one state, it is assigned as being dynamic. (E) Dwell-based 2D E-S scatter plot. Dwells are colored on the basis of the assigned state of the chosen mpH2MM model. The dwells were corrected for leakage, direct excitation and the γ-factor. Black dots in A, D and E represent the average value of each state.

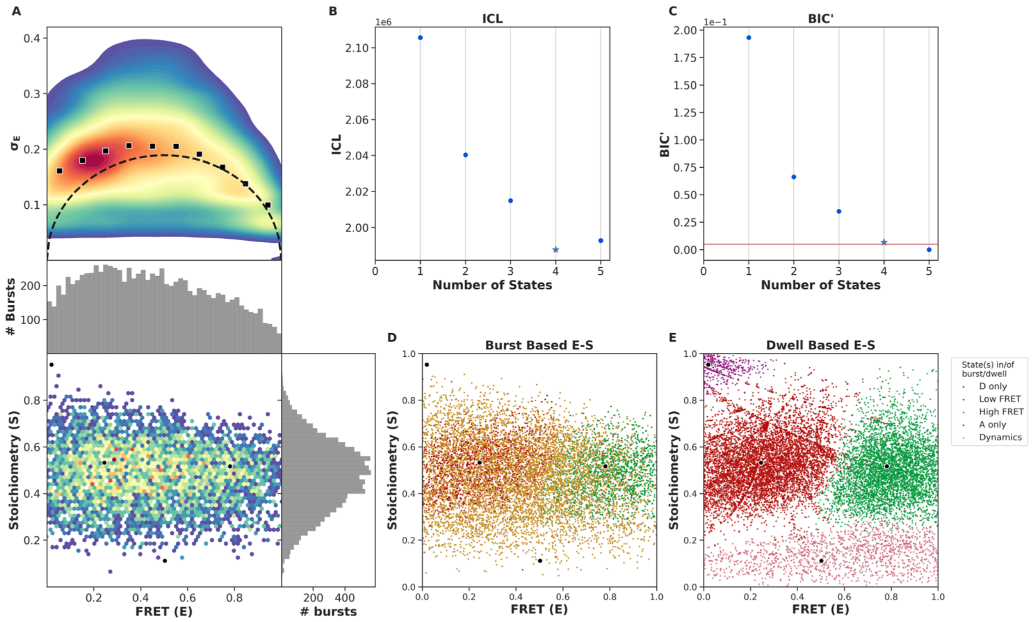


FIGURE S3. Burst variance analysis (BVA) and mpH2MM of the FRET bursts of OpuA-K521C in 50 mM HEPES-K pH 7.0 plus 100 µM glycine betaine. (A) From top to bottom: (1) Burst variance analysis of the bursts which were corrected by the leakage, crosstalk, and γ-correction factors and which were selected after removing donor-only and acceptor-only bursts. The standard deviation of FRET in each burst is plotted against its mean FRET. Black squares show average values per FRET bin. Black dotted line shows the expected standard deviation in the absence of within-burst dynamics. (2) 2D E-S histogram shows the same data as in (1), with on both sides a histogram that represents the same bursts. (B) Plot of the ICL-values for each final model. The model used in the downstream analysis and following figures is shown as a star. (C) Plot of the BIC'-values for each final model. The red line represents a 0.05 cut-off. The model used is shown as a star. (D) Burst-based 2D E-S scatter plot. Bursts are colored on the basis of the assigned state of the chosen mpH2MM model. If a burst contains more than one state, it is assigned as being dynamic. (E) Dwell-based 2D E-S scatter plot. Dwells are colored on the basis of the assigned state of the chosen mpH2MM model. The dwells were corrected for leakage, direct excitation and the γ-factor. Black dots in A, D and E represent the average value of each state.

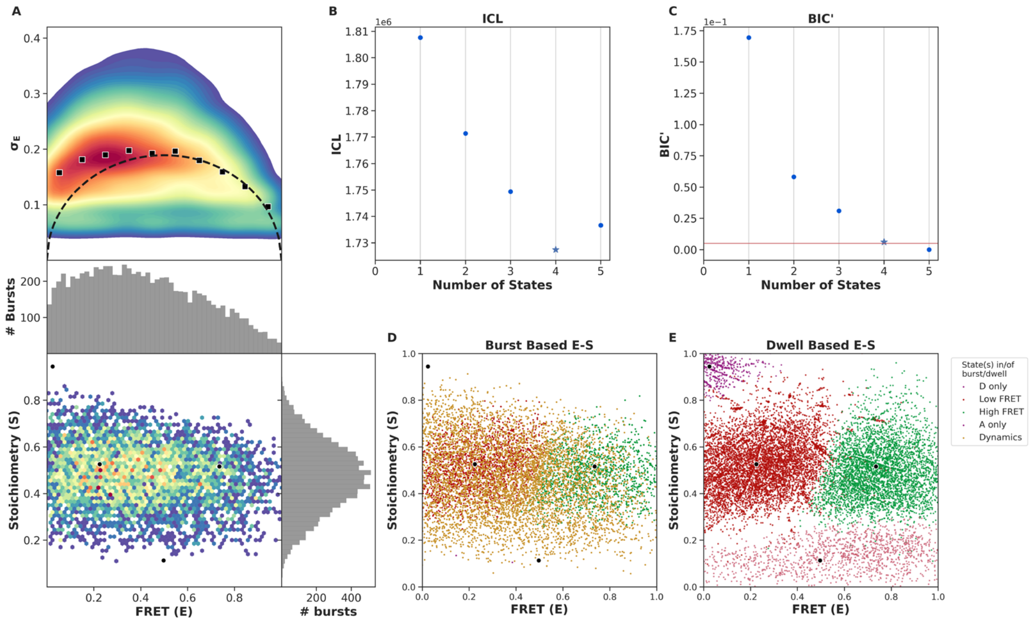


FIGURE S4. Burst variance analysis (BVA) and mpH2MM of the FRET bursts of OpuA-K521C in 50 mM HEPES-K pH 7.0 plus 50 mM KCl. (A) From top to bottom: (1) Burst variance analysis of the bursts which were corrected by the leakage, crosstalk, and γ-correction factors and which were selected after removing donor-only and acceptor-only bursts. The standard deviation of FRET in each burst is plotted against its mean FRET. Black squares show average values per FRET bin. Black dotted line shows the expected standard deviation in the absence of within-burst dynamics. (2) 2D E-S histogram shows the same data as in (1), with on both sides a histogram that represents the same bursts. (B) Plot of the ICL-values for each final model. The model used in the downstream analysis and following figures is shown as a star. (C) Plot of the BIC'-values for each final model. The red line represents a 0.05 cut-off. The model used is shown as a star. (D) Burst-based 2D E-S scatter plot. Bursts are colored on the basis of the assigned state of the chosen mpH2MM model. If a burst contains more than one state, it is assigned as being dynamic. (E) Dwell-based 2D E-S scatter plot. Dwells are colored on the basis of the assigned state of the chosen mpH2MM model. The dwells were corrected for leakage, direct excitation and the γ-factor. Black dots in A, D and E represent the average value of each state.

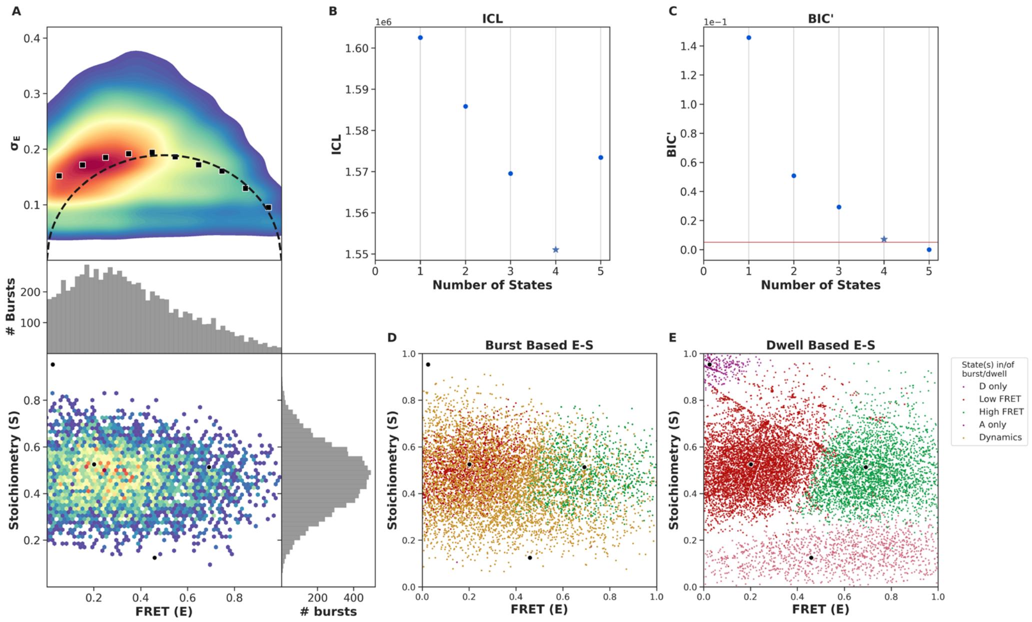


FIGURE S5. Burst variance analysis (BVA) and mpH2MM of the FRET bursts of OpuA-K521C in 50 mM HEPES-K pH 7.0 plus 100 mM KCl. (A) From top to bottom: (1) Burst variance analysis of the bursts which were corrected by the leakage, crosstalk, and γ-correction factors and which were selected after removing donor-only and acceptor-only bursts. The standard deviation of FRET in each burst is plotted against its mean FRET. Black squares show average values per FRET bin. Black dotted line shows the expected standard deviation in the absence of within-burst dynamics. (2) 2D E-S histogram shows the same data as in (1), with on both sides a histogram that represents the same bursts. (B) Plot of the ICL-values for each final model. The model used in the downstream analysis and following figures is shown as a star. (C) Plot of the BIC'-values for each final model. The red line represents a 0.05 cut-off. The model used is shown as a star. (D) Burst-based 2D E-S scatter plot. Bursts are colored on the basis of the assigned state of the chosen mpH2MM model. If a burst contains more than one state, it is assigned as being dynamic. (E) Dwell-based 2D E-S scatter plot. Dwells are colored on the basis of the assigned state of the chosen mpH2MM model. The dwells were corrected for leakage, direct excitation and the γ-factor. Black dots in A, D and E represent the average value of each state..

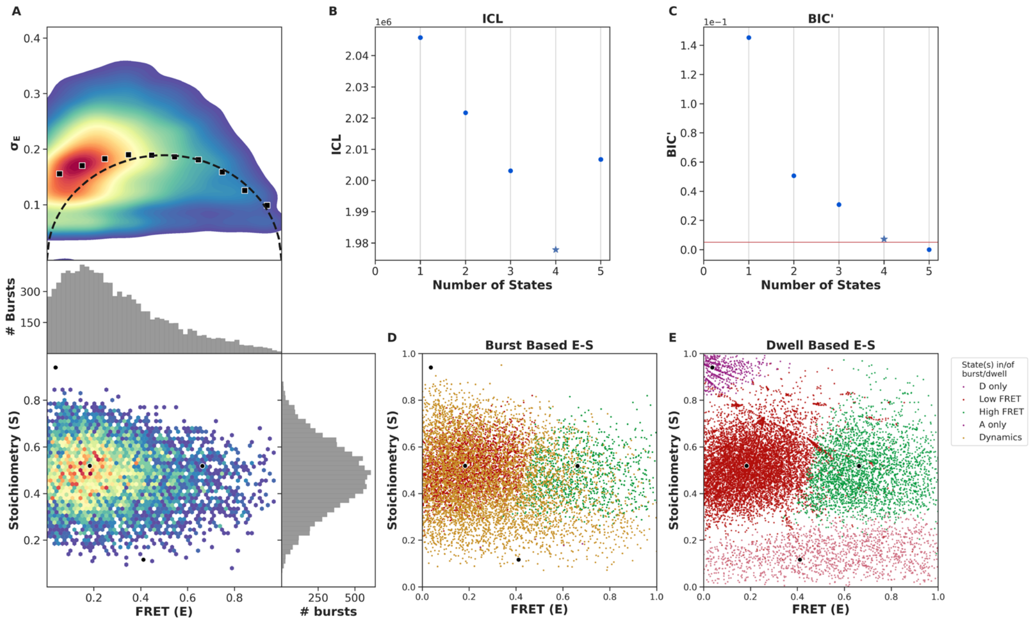


FIGURE S6. Burst variance analysis (BVA) and mpH2MM of the FRET bursts of OpuA-K521C in 50 mM HEPES-K pH 7.0 plus 200 mM KCl. (A) From top to bottom: (1) Burst variance analysis of the bursts which were corrected by the leakage, crosstalk, and γ-correction factors and which were selected after removing donor-only and acceptor-only bursts. The standard deviation of FRET in each burst is plotted against its mean FRET. Black squares show average values per FRET bin. Black dotted line shows the expected standard deviation in the absence of within-burst dynamics. (2) 2D E-S histogram shows the same data as in (1), with on both sides a histogram that represents the same bursts. (B) Plot of the ICL-values for each final model. The model used in the downstream analysis and following figures is shown as a star. (C) Plot of the BIC'-values for each final model. The red line represents a 0.05 cut-off. The model used is shown as a star. (D) Burst-based 2D E-S scatter plot. Bursts are colored on the basis of the assigned state of the chosen mpH2MM model. If a burst contains more than one state, it is assigned as being dynamic. (E) Dwell-based 2D E-S scatter plot. Dwells are colored on the basis of the assigned state of the chosen mpH2MM model. The dwells were corrected for leakage, direct excitation and the γ-factor. Black dots in A, D and E represent the average value of each state.

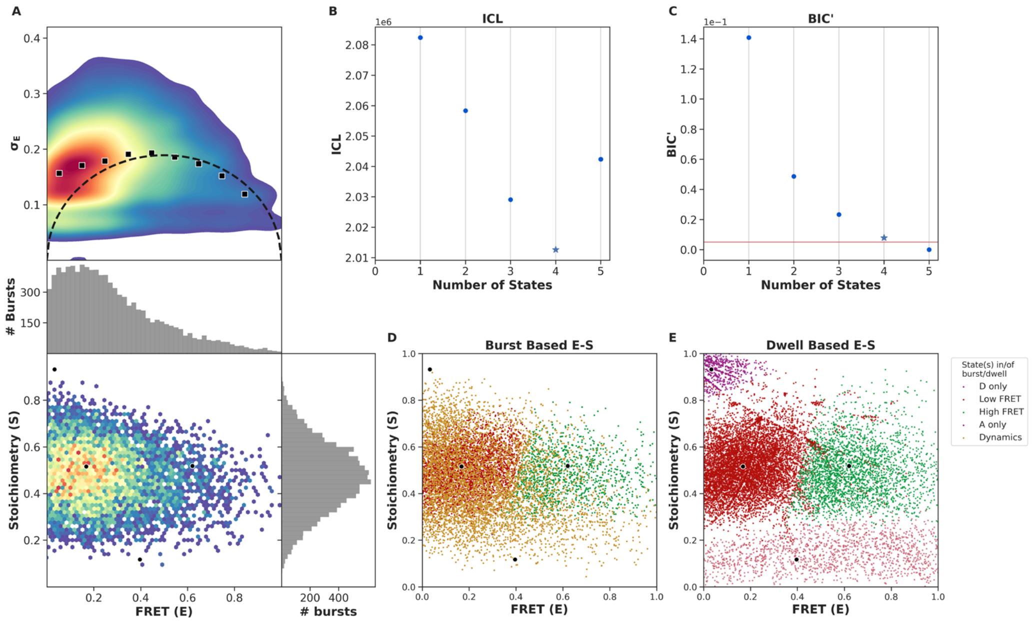


FIGURE S7. Burst variance analysis (BVA) and mpH2MM of the FRET bursts of OpuA-K521C in 50 mM HEPES-K pH 7.0 plus 400 mM KCl. (A) From top to bottom: (1) Burst variance analysis of the bursts which were corrected by the leakage, crosstalk, and γ-correction factors and which were selected after removing donor-only and acceptor-only bursts. The standard deviation of FRET in each burst is plotted against its mean FRET. Black squares show average values per FRET bin. Black dotted line shows the expected standard deviation in the absence of within-burst dynamics. (2) 2D E-S histogram shows the same data as in (1), with on both sides a histogram that represents the same bursts. (B) Plot of the ICL-values for each final model. The model used in the downstream analysis and following figures is shown as a star. (C) Plot of the BIC'-values for each final model. The red line represents a 0.05 cut-off. The model used is shown as a star. (D) Burst-based 2D E-S scatter plot. Bursts are colored on the basis of the assigned state of the chosen mpH2MM model. If a burst contains more than one state, it is assigned as being dynamic. (E) Dwell-based 2D E-S scatter plot. Dwells are colored on the basis of the assigned state of the chosen mpH2MM model. The dwells were corrected for leakage, direct excitation and the γ-factor. Black dots in A, D and E represent the average value of each state.

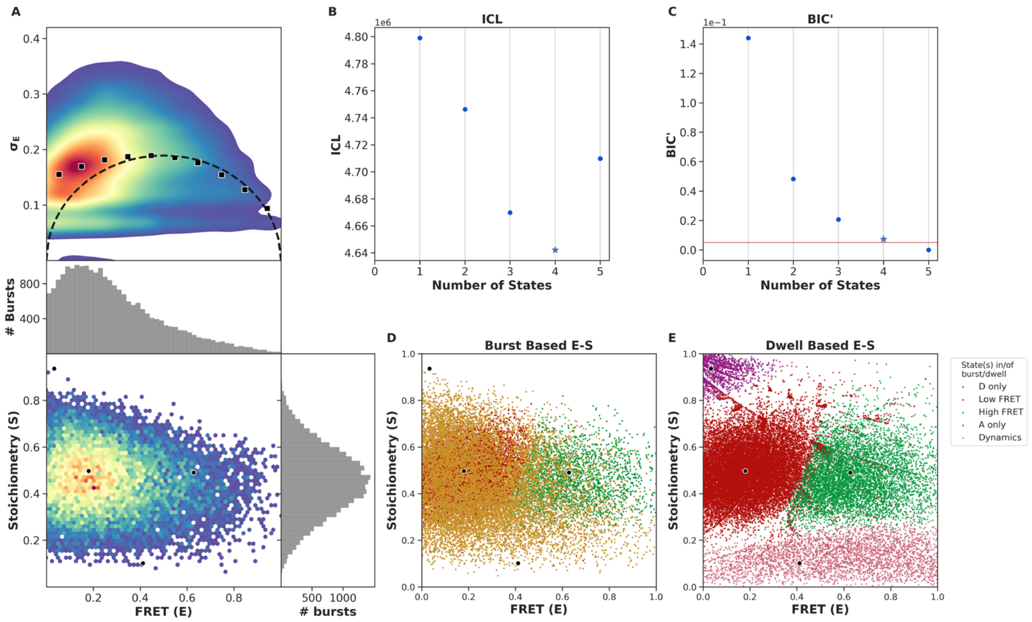


FIGURE S8. Burst variance analysis (BVA) and mpH2MM of the FRET bursts of OpuA-K521C in 50 mM HEPES-K pH 7.0 plus 600 mM KCl. (A) From top to bottom: (1) Burst variance analysis of the bursts which were corrected by the leakage, crosstalk, and γ-correction factors and which were selected after removing donor-only and acceptor-only bursts. The standard deviation of FRET in each burst is plotted against its mean FRET. Black squares show average values per FRET bin. Black dotted line shows the expected standard deviation in the absence of within-burst dynamics. (2) 2D E-S histogram shows the same data as in (1), with on both sides a histogram that represents the same bursts. (B) Plot of the ICL-values for each final model. The model used in the downstream analysis and following figures is shown as a star. (C) Plot of the BIC'-values for each final model. The red line represents a 0.05 cut-off. The model used is shown as a star. (D) Burst-based 2D E-S scatter plot. Bursts are colored on the basis of the assigned state of the chosen mpH2MM model. If a burst contains more than one state, it is assigned as being dynamic. (E) Dwell-based 2D E-S scatter plot. Dwells are colored on the basis of the assigned state of the chosen mpH2MM model. The dwells were corrected for leakage, direct excitation and the γ-factor. Black dots in A, D and E represent the average value of each state.

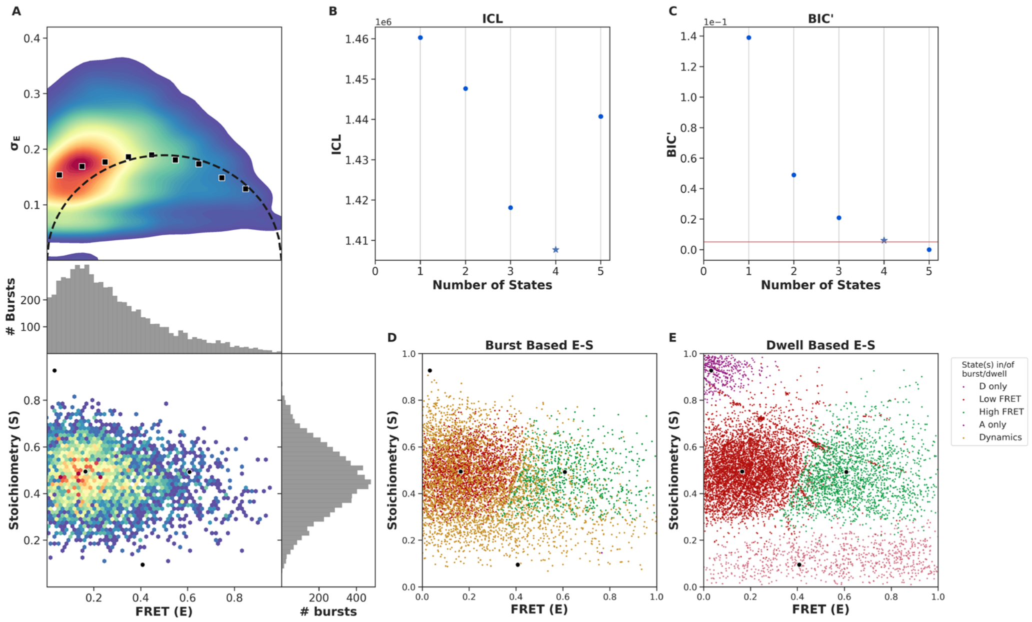


FIGURE S9. Burst variance analysis (BVA) and mpH2MM of the FRET bursts of OpuA-V149Q-K521C in 50 mM HEPES-K pH 7.0 plus 600 mM KCl. (A) From top to bottom: (1) Burst variance analysis of the bursts which were corrected by the leakage, crosstalk, and γ-correction factors and which were selected after removing donor-only and acceptor-only bursts. The standard deviation of FRET in each burst is plotted against its mean FRET. Black squares show average values per FRET bin. Black dotted line shows the expected standard deviation in the absence of within-burst dynamics. (2) 2D E-S histogram shows the same data as in (1), with on both sides a histogram that represents the same bursts. (B) Plot of the ICL-values for each final model. The model used in the downstream analysis and following figures is shown as a star. (C) Plot of the BIC'-values for each final model. The red line represents a 0.05 cut-off. The model used is shown as a star. (D) Burst-based 2D E-S scatter plot. Bursts are colored on the basis of the assigned state of the chosen mpH2MM model. If a burst contains more than one state, it is assigned as being dynamic. (E) Dwell-based 2D E-S scatter plot. Dwells are colored on the basis of the assigned state of the chosen mpH2MM model. The dwells were corrected for leakage, direct excitation and the γ-factor. Black dots in A, D and E represent the average value of each state.

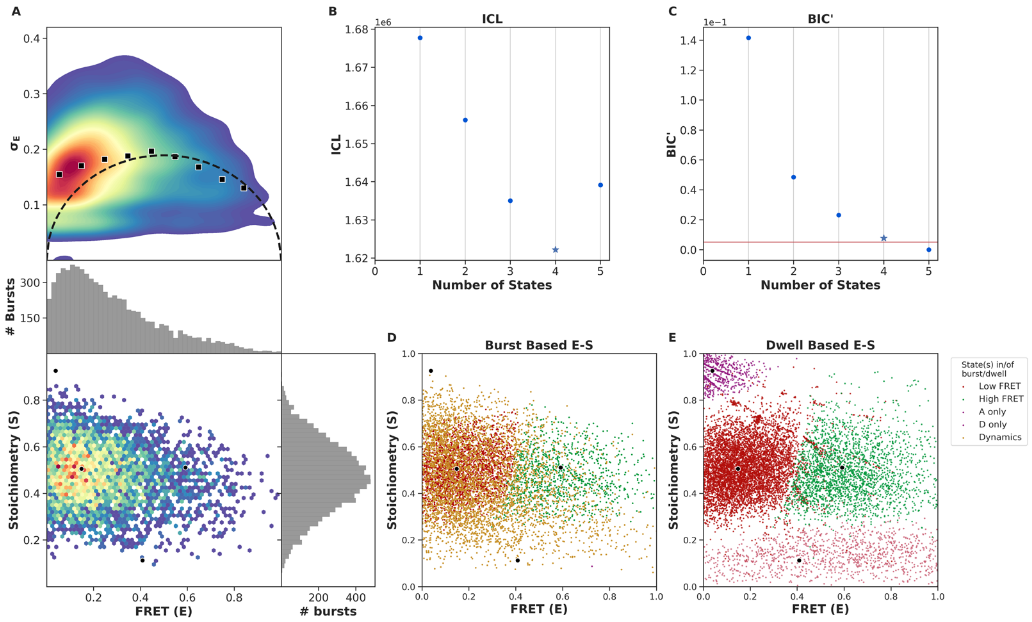


FIGURE S10. Burst variance analysis (BVA) and mpH2MM of the FRET bursts of OpuA-K521C in 50 mM HEPES-K pH 7.0 plus 1000 mM KCl. (A) From top to bottom: (1) Burst variance analysis of the bursts which were corrected by the leakage, crosstalk, and γ-correction factors and which were selected after removing donor-only and acceptor-only bursts. The standard deviation of FRET in each burst is plotted against its mean FRET. Black squares show average values per FRET bin. Black dotted line shows the expected standard deviation in the absence of within-burst dynamics. (2) 2D E-S histogram shows the same data as in (1), with on both sides a histogram that represents the same bursts. (B) Plot of the ICL-values for each final model. The model used in the downstream analysis and following figures is shown as a star. (C) Plot of the BIC'-values for each final model. The red line represents a 0.05 cut-off. The model used is shown as a star. (D) Burst-based 2D E-S scatter plot. Bursts are colored on the basis of the assigned state of the chosen mpH2MM model. If a burst contains more than one state, it is assigned as being dynamic. (E) Dwell-based 2D E-S scatter plot. Dwells are colored on the basis of the assigned state of the chosen mpH2MM model. The dwells were corrected for leakage, direct excitation and the γ-factor. Black dots in A, D and E represent the average value of each state.

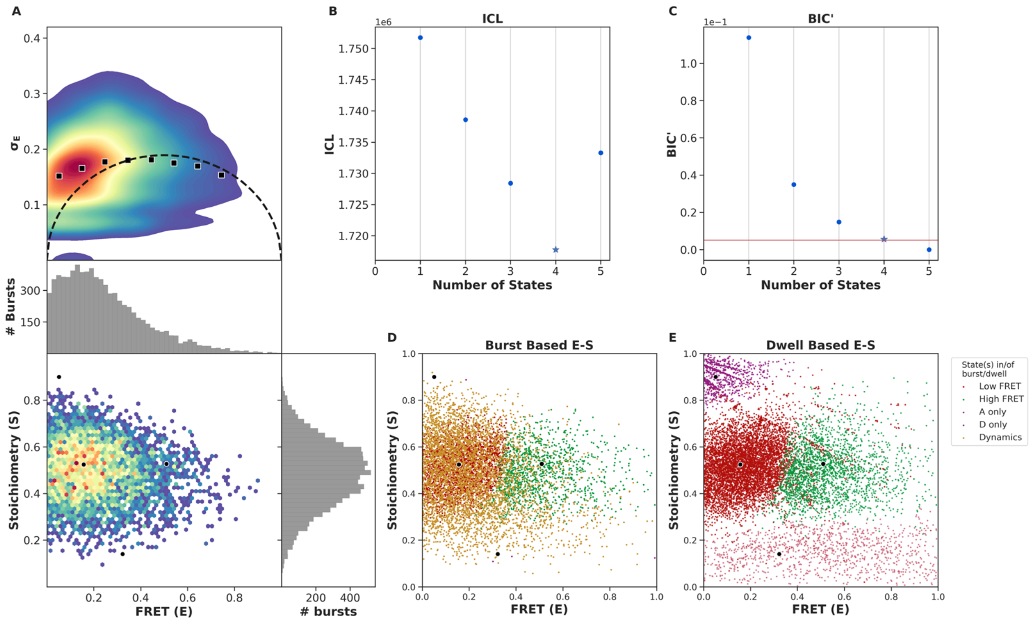


FIGURE S11. Burst variance analysis (BVA) and mpH2MM of the FRET bursts of OpuA-K521C in 50 mM HEPES-K pH 7.0, 600 mM KCl, 20 mM Mg-ATP. (A) From top to bottom: (1) Burst variance analysis of the bursts which were corrected by the leakage, crosstalk, and γ-correction factors and which were selected after removing donor-only and acceptor-only bursts. The standard deviation of FRET in each burst is plotted against its mean FRET. Black squares show average values per FRET bin. Black dotted line shows the expected standard deviation in the absence of within-burst dynamics. (2) 2D E-S histogram shows the same data as in (1), with on both sides a histogram that represents the same bursts. (B) Plot of the ICL-values for each final model. The model used in the downstream analysis and following figures is shown as a star. (C) Plot of the BIC'-values for each final model. The red line represents a 0.05 cut-off. The model used is shown as a star. (D) Burst-based 2D E-S scatter plot. Bursts are colored on the basis of the assigned state of the chosen mpH2MM model. If a burst contains more than one state, it is assigned as being dynamic. (E) Dwell-based 2D E-S scatter plot. Dwells are colored on the basis of the assigned state of the chosen mpH2MM model. The dwells were corrected for leakage, direct excitation and the γ-factor. Black dots in A, D and E represent the average value of each state.

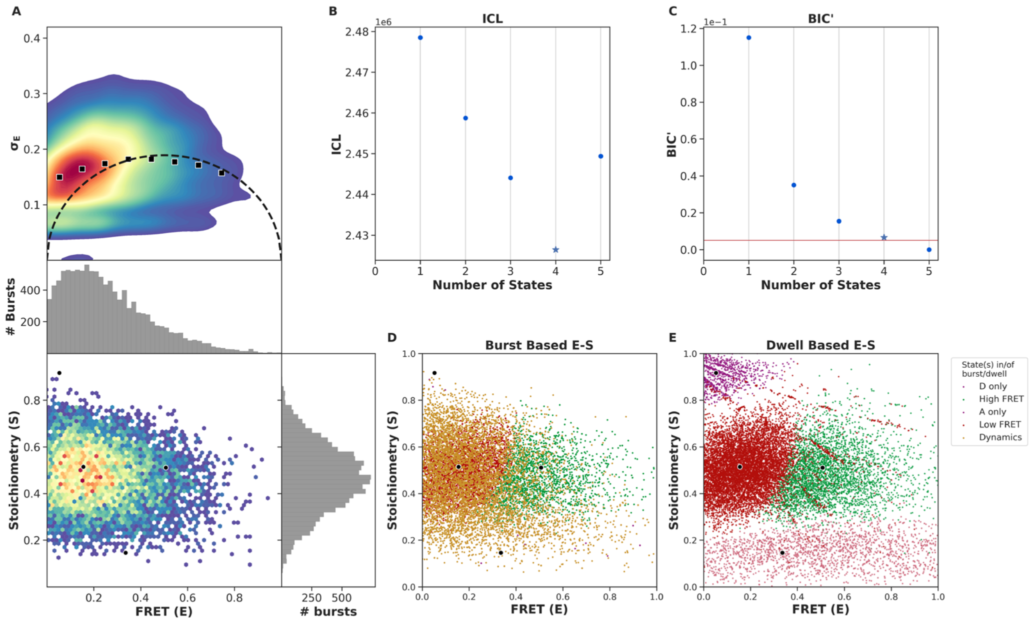


FIGURE S12. Burst variance analysis (BVA) and mpH2MM of the FRET bursts of OpuA-K521C in 50 mM HEPES-K pH 7.0, 600 mM KCl, 20 mM Mg-ATP plus 100 µM glycine betaine. (A) From top to bottom: (1) Burst variance analysis of the bursts which were corrected by the leakage, crosstalk, and γ-correction factors and which were selected after removing donor-only and acceptor-only bursts. The standard deviation of FRET in each burst is plotted against its mean FRET. Black squares show average values per FRET bin. Black dotted line shows the expected standard deviation in the absence of within-burst dynamics. (2) 2D E-S histogram shows the same data as in (1), with on both sides a histogram that represents the same bursts. (B) Plot of the ICL-values for each final model. The model used in the downstream analysis and following figures is shown as a star. (C) Plot of the BIC'-values for each final model. The red line represents a 0.05 cut-off. The model used is shown as a star. (D) Burst-based 2D E-S scatter plot. Bursts are colored on the basis of the assigned state of the chosen mpH2MM model. If a burst contains more than one state, it is assigned as being dynamic. (E) Dwell-based 2D E-S scatter plot. Dwells are colored on the basis of the assigned state of the chosen mpH2MM model. The dwells were corrected for leakage, direct excitation and the γ-factor. Black dots in A, D and E represent the average value of each state.

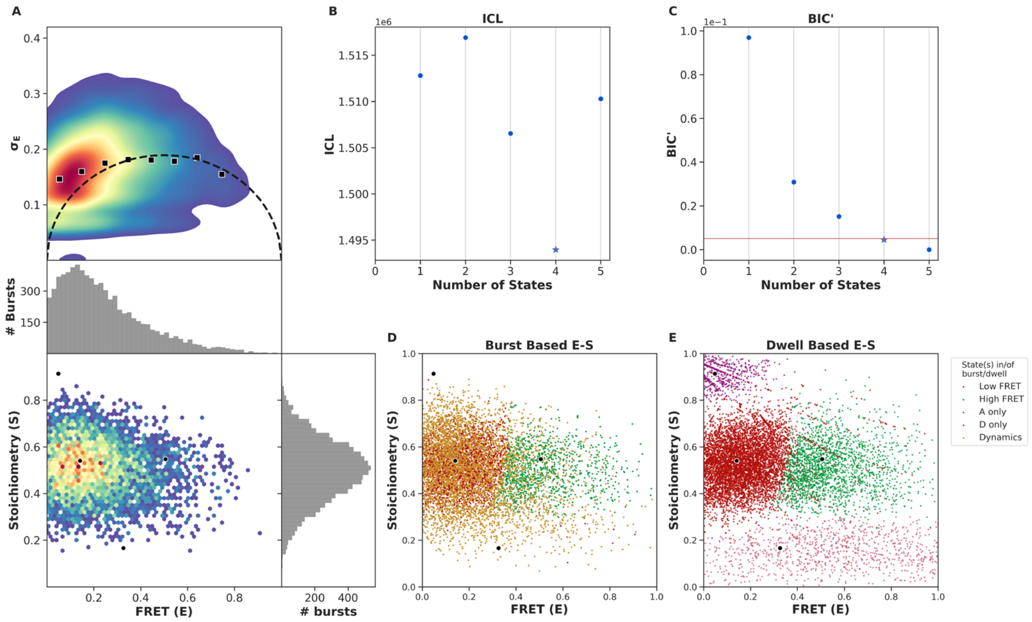


FIGURE S13. Burst variance analysis (BVA) and mpH2MM of the FRET bursts of OpuA-E190Q-K521C in 50 mM HEPES-K pH 7.0, 600 mM KCl, 20 mM Mg-ATP plus 100 μM glycine betaine. (A) From top to bottom: (1) Burst variance analysis of the bursts which were corrected by the leakage, crosstalk, and γ-correction factors and which were selected after removing donor-only and acceptor-only bursts. The standard deviation of FRET in each burst is plotted against its mean FRET. Black squares show average values per FRET bin. Black dotted line shows the expected standard deviation in the absence of within-burst dynamics. (2) 2D E-S histogram shows the same data as in (1), with on both sides a histogram that represents the same bursts. (B) Plot of the ICL-values for each final model. The model used in the downstream analysis and following figures is shown as a star. (C) Plot of the BIC'-values for each final model. The red line represents a 0.05 cut-off. The model used is shown as a star. (D) Burst-based 2D E-S scatter plot. Bursts are colored on the basis of the assigned state of the chosen mpH2MM model. If a burst contains more than one state, it is assigned as being dynamic. (E) Dwell-based 2D E-S scatter plot. Dwells are colored on the basis of the assigned state of the chosen mpH2MM model. The dwells were corrected for leakage, direct excitation and the γ-factor. Black dots in A, D and E represent the average value of each state.

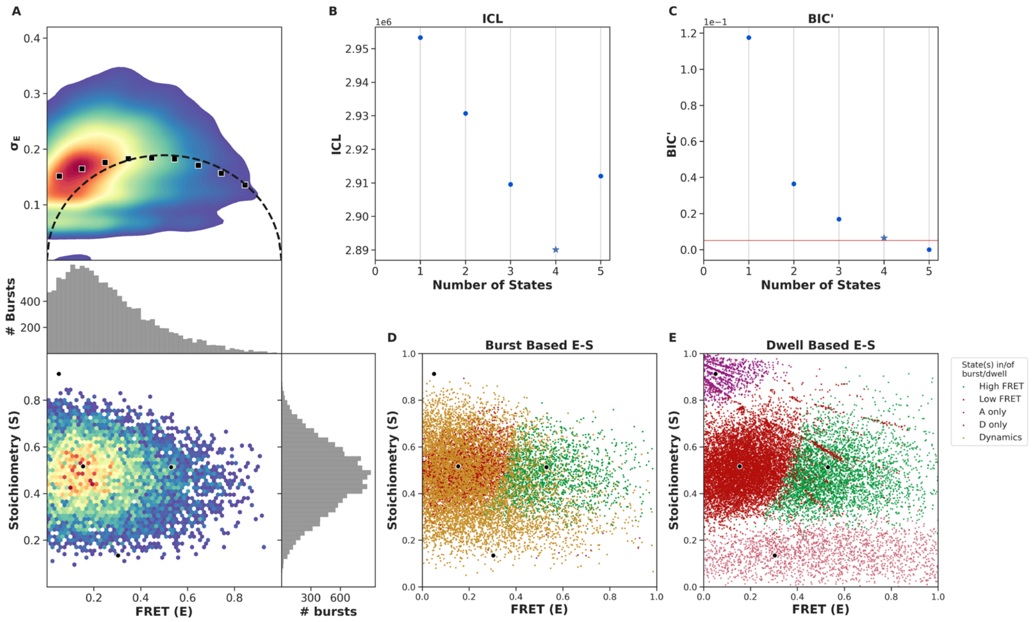


FIGURE S14. Burst variance analysis (BVA) and mpH2MM of the FRET bursts of OpuA-K521C in 50 mM HEPES-K pH 7.0, 600 mM KCl, 20 mM Mg-ATP, 100 µM glycine betaine plus 500 µM *ortho*vanadate. (A) From top to bottom: (1) Burst variance analysis of the bursts which were corrected by the leakage, crosstalk, and γ-correction factors and which were selected after removing donor-only and acceptor-only bursts. The standard deviation of FRET in each burst is plotted against its mean FRET. Black squares show average values per FRET bin. Black dotted line shows the expected standard deviation in the absence of within-burst dynamics. (2) 2D E-S histogram shows the same data as in (1), with on both sides a histogram that represents the same bursts. (B) Plot of the ICL-values for each final model. The model used in the downstream analysis and following figures is shown as a star. (C) Plot of the BIC'-values for each final model. The red line represents a 0.05 cut-off. The model used is shown as a star. (D) Burst-based 2D E-S scatter plot. Bursts are colored on the basis of the assigned state of the chosen mpH2MM model. If a burst contains more than one state, it is assigned as being dynamic. (E) Dwell-based 2D E-S scatter plot. Dwells are colored on the basis of the assigned state of the chosen mpH2MM model. The dwells were corrected for leakage, direct excitation and the γ-factor. Black dots in A, D and E represent the average value of each state.

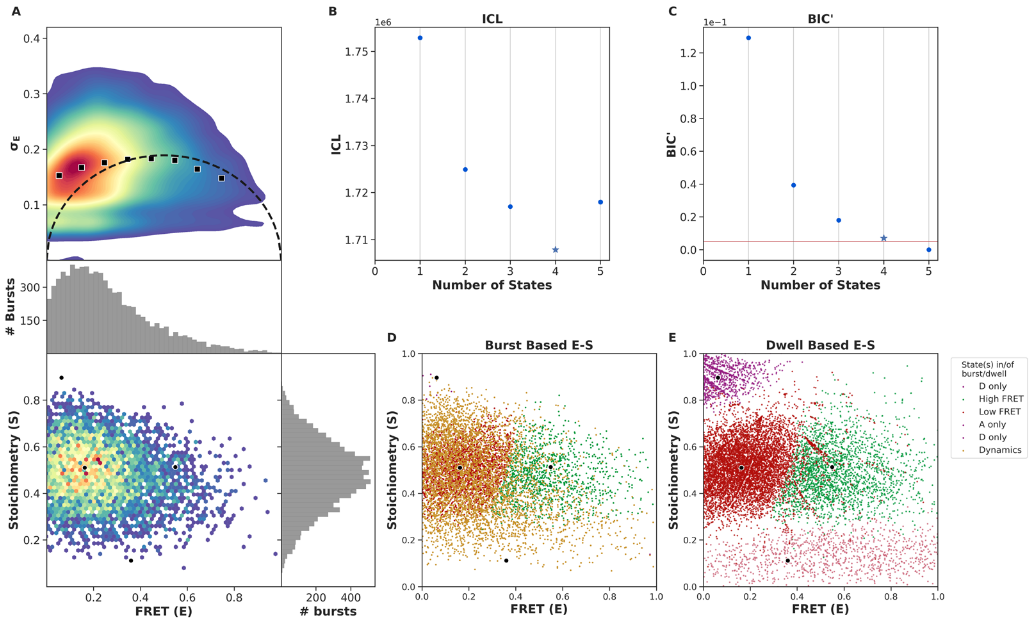


FIGURE S15. Burst variance analysis (BVA) and mpH2MM of the FRET bursts of OpuA-K521C in 50 mM HEPES-K pH 7.0, 600 mM KCl, 50 mM arginine plus 50 mM glutamate. (A) From top to bottom: (1) Burst variance analysis of the bursts which were corrected by the leakage, crosstalk, and γ-correction factors and which were selected after removing donor-only and acceptor-only bursts. The standard deviation of FRET in each burst is plotted against its mean FRET. Black squares show average values per FRET bin. Black dotted line shows the expected standard deviation in the absence of within-burst dynamics. (2) 2D E-S histogram shows the same data as in (1), with on both sides a histogram that represents the same bursts. (B) Plot of the ICL-values for each final model. The model used in the downstream analysis and following figures is shown as a star. (C) Plot of the BIC'-values for each final model. The red line represents a 0.05 cut-off. The model used is shown as a star. (D) Burst-based 2D E-S scatter plot. Bursts are colored on the basis of the assigned state of the chosen mpH2MM model. If a burst contains more than one state, it is assigned as being dynamic. (E) Dwell-based 2D E-S scatter plot. Dwells are colored on the basis of the assigned state of the chosen mpH2MM model. The dwells were corrected for leakage, direct excitation and the γ-factor. Black dots in A, D and E represent the average value of each state.

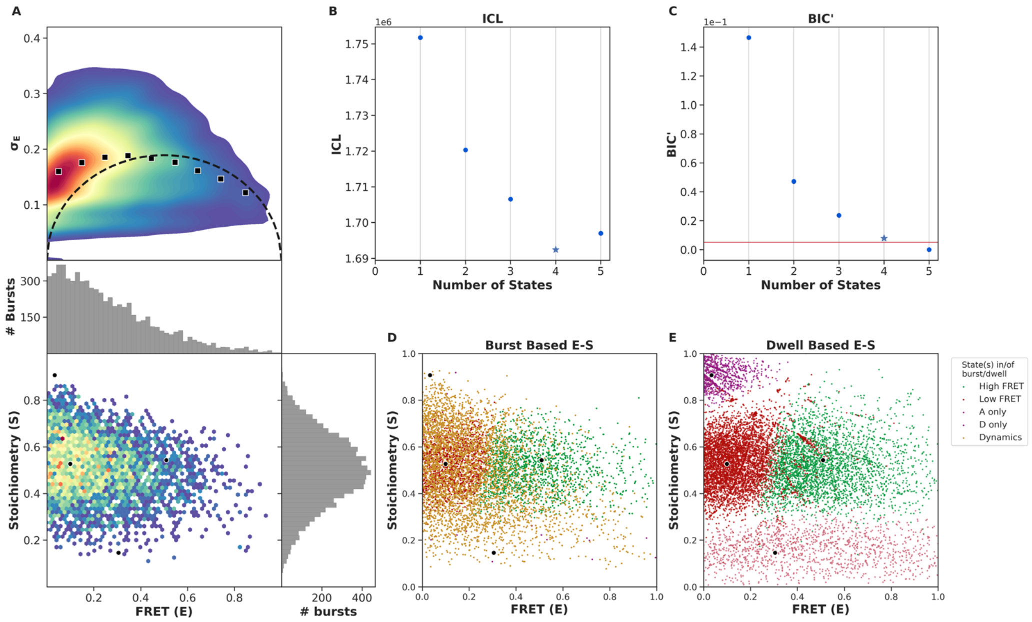


FIGURE S16. Burst variance analysis (BVA) and mpH2MM of the FRET bursts of OpuA-N414C in 50 mM HEPES-K pH 7.0. (A) From top to bottom: (1) Burst variance analysis of the bursts which were corrected by the leakage, crosstalk, and γ-correction factors and which were selected after removing donor-only and acceptor-only bursts. The standard deviation of FRET in each burst is plotted against its mean FRET. Black squares show average values per FRET bin. Black dotted line shows the expected standard deviation in the absence of within-burst dynamics. (2) 2D E-S histogram shows the same data as in (1), with on both sides a histogram that represents the same bursts. (B) Plot of the ICL-values for each final model. The model used in the downstream analysis and following figures is shown as a star. (C) Plot of the BIC'-values for each final model. The red line represents a 0.05 cut-off. The model used is shown as a star. (D) Burst-based 2D E-S scatter plot. Bursts are colored on the basis of the assigned state of the chosen mpH2MM model. If a burst contains more than one state, it is assigned as being dynamic. (E) Dwell-based 2D E-S scatter plot. Dwells are colored on the basis of the assigned state of the chosen mpH2MM model. The dwells were corrected for leakage, direct excitation and the γ-factor. Black dots in A, D and E represent the average value of each state.

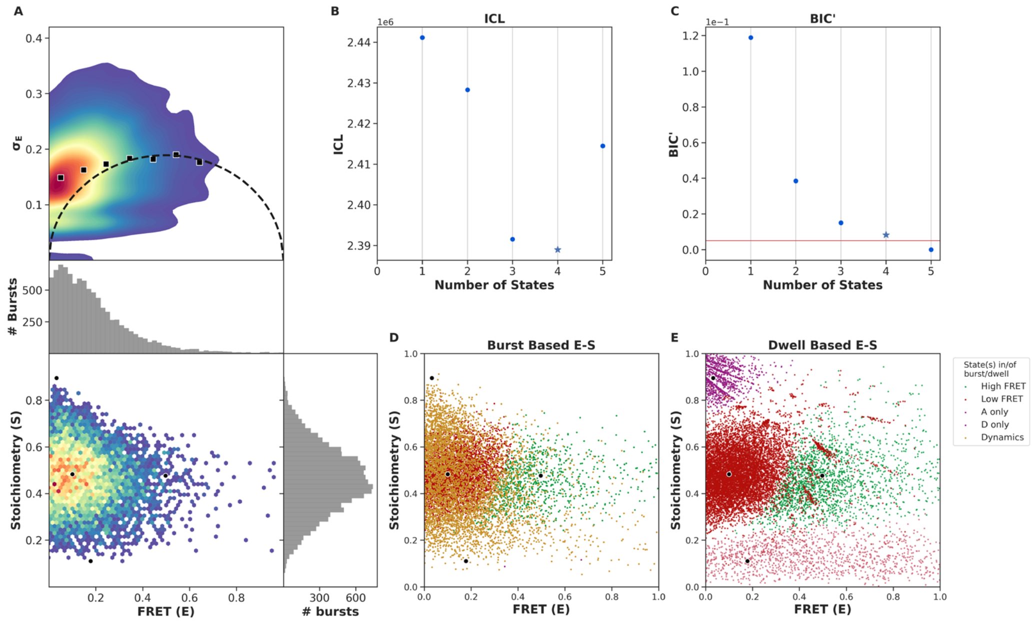


FIGURE S17. Burst variance analysis (BVA) and mpH2MM of the FRET bursts of OpuA-N414C in 50 mM HEPES-K pH 7.0 plus 600 mM KCl. (A) From top to bottom: (1) Burst variance analysis of the bursts which were corrected by the leakage, crosstalk, and γ-correction factors and which were selected after removing donor-only and acceptor-only bursts. The standard deviation of FRET in each burst is plotted against its mean FRET. Black squares show average values per FRET bin. Black dotted line shows the expected standard deviation in the absence of within-burst dynamics. (2) 2D E-S histogram shows the same data as in (1), with on both sides a histogram that represents the same bursts. (B) Plot of the ICL-values for each final model. The model used in the downstream analysis and following figures is shown as a star. (C) Plot of the BIC'-values for each final model. The red line represents a 0.05 cut-off. The model used is shown as a star. (D) Burst-based 2D E-S scatter plot. Bursts are colored on the basis of the assigned state of the chosen mpH2MM model. If a burst contains more than one state, it is assigned as being dynamic. (E) Dwell-based 2D E-S scatter plot. Dwells are colored on the basis of the assigned state of the chosen mpH2MM model. The dwells were corrected for leakage, direct excitation and the γ-factor. Black dots in A, D and E represent the average value of each state.

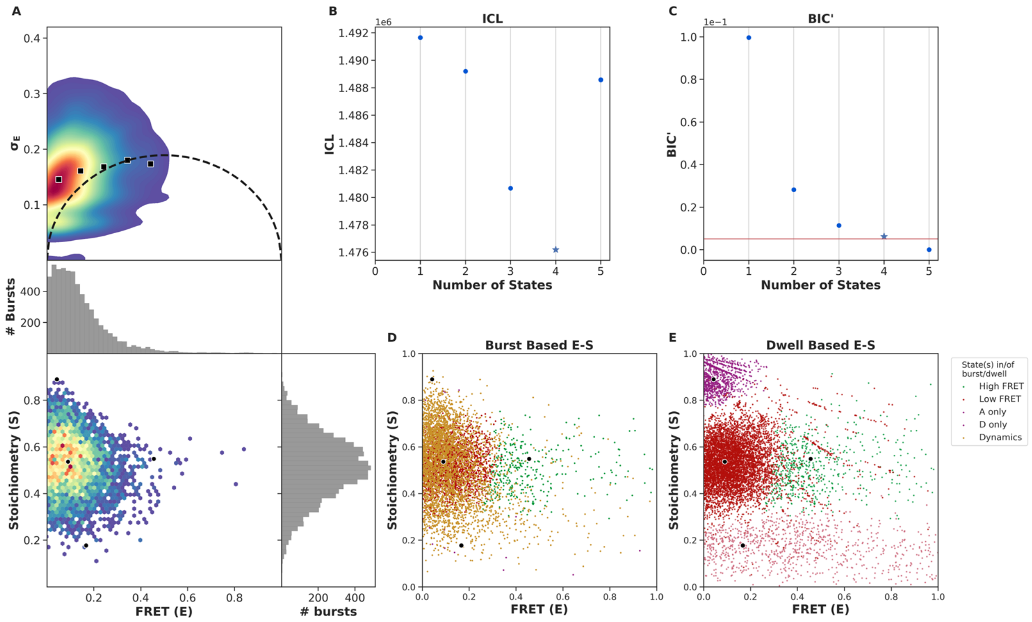


FIGURE S18. Burst variance analysis (BVA) and mpH2MM of the FRET bursts of OpuA-N414C in 50 mM HEPES-K pH 7.0, 600 mM KCl plus 20 mM Mg-ATP. (A) From top to bottom: (1) Burst variance analysis of the bursts which were corrected by the leakage, crosstalk, and γ-correction factors and which were selected after removing donor-only and acceptor-only bursts. The standard deviation of FRET in each burst is plotted against its mean FRET. Black squares show average values per FRET bin. Black dotted line shows the expected standard deviation in the absence of within-burst dynamics. (2) 2D E-S histogram shows the same data as in (1), with on both sides a histogram that represents the same bursts. (B) Plot of the ICL-values for each final model. The model used in the downstream analysis and following figures is shown as a star. (C) Plot of the BIC'-values for each final model. The red line represents a 0.05 cut-off. The model used is shown as a star. (D) Burst-based 2D E-S scatter plot. Bursts are colored on the basis of the assigned state of the chosen mpH2MM model. If a burst contains more than one state, it is assigned as being dynamic. (E) Dwell-based 2D E-S scatter plot. Dwells are colored on the basis of the assigned state of the chosen mpH2MM model. The dwells were corrected for leakage, direct excitation and the γ-factor. Black dots in A, D and E represent the average value of each state.

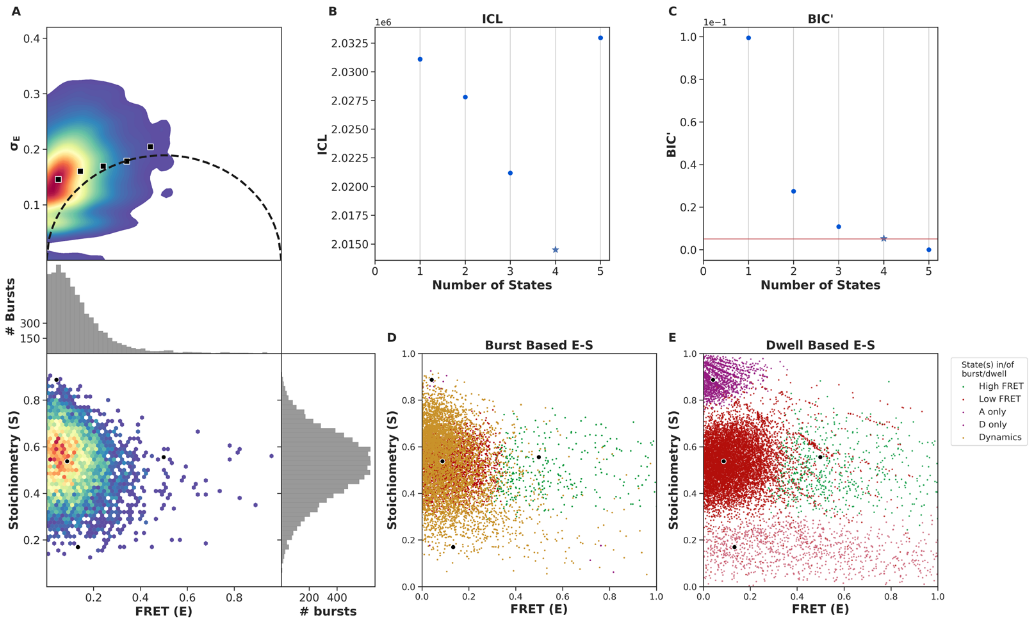


FIGURE S19. Burst variance analysis (BVA) and mpH2MM of the FRET bursts of OpuA-N414C in 50 mM HEPES-K pH 7.0, 600 mM KCl, 20 mM Mg-ATP plus 100 µM glycine betaine. (A) From top to bottom: (1) Burst variance analysis of the bursts which were corrected by the leakage, crosstalk, and γ-correction factors and which were selected after removing donor-only and acceptor-only bursts. The standard deviation of FRET in each burst is plotted against its mean FRET. Black squares show average values per FRET bin. Black dotted line shows the expected standard deviation in the absence of within-burst dynamics. (2) 2D E-S histogram shows the same data as in (1), with on both sides a histogram that represents the same bursts. (B) Plot of the ICL-values for each final model. The model used in the downstream analysis and following figures is shown as a star. (C) Plot of the BIC'-values for each final model. The red line represents a 0.05 cut-off. The model used is shown as a star. (D) Burst-based 2D E-S scatter plot. Bursts are colored on the basis of the assigned state of the chosen mpH2MM model. If a burst contains more than one state, it is assigned as being dynamic. (E) Dwell-based 2D E-S scatter plot. Dwells are colored on the basis of the assigned state of the chosen mpH2MM model. The dwells were corrected for leakage, direct excitation and the γ-factor. Black dots in A, D and E represent the average value of each state.

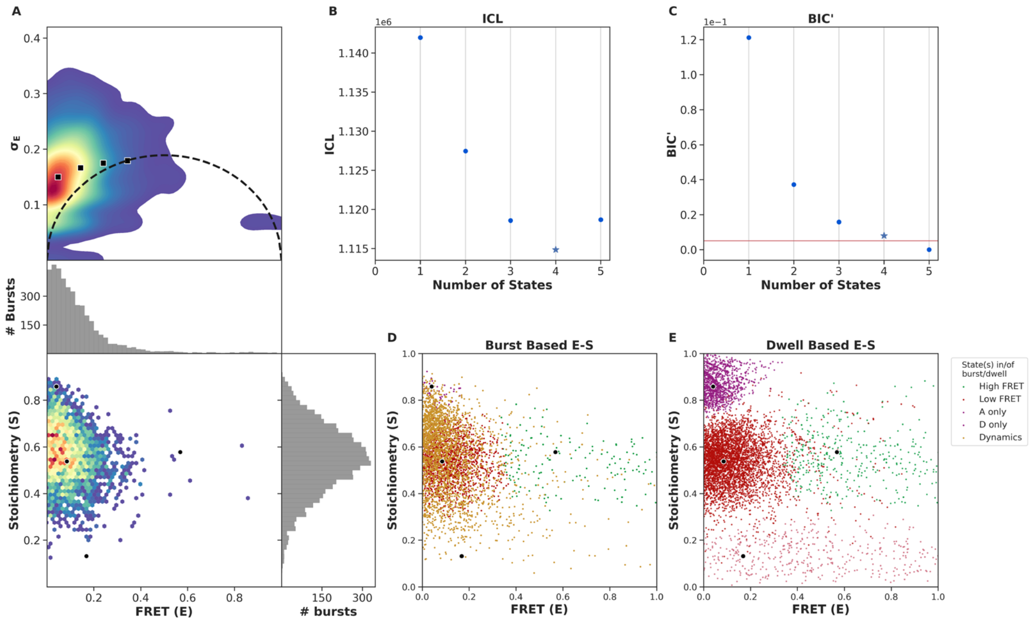


FIGURE S20. Burst variance analysis (BVA) and mpH2MM of the FRET bursts of OpuA-N414C in 50 mM HEPES-K pH 7.0, 600 mM KCl, 20 mM Mg-ATP, 100 µM glycine betaine plus 500 µM *ortho*vanadate. (A) From top to bottom: (1) Burst variance analysis of the bursts which were corrected by the leakage, crosstalk, and γ-correction factors and which were selected after removing donor-only and acceptor-only bursts. The standard deviation of FRET in each burst is plotted against its mean FRET. Black squares show average values per FRET bin. Black dotted line shows the expected standard deviation in the absence of within-burst dynamics. (2) 2D E-S histogram shows the same data as in (1), with on both sides a histogram that represents the same bursts. (B) Plot of the ICL-values for each final model. The model used in the downstream analysis and following figures is shown as a star. (C) Plot of the BIC’-values for each final model. The red line represents a 0.05 cut-off. The model used is shown as a star. (D) Burst-based 2D E-S scatter plot. Bursts are colored on the basis of the assigned state of the chosen mpH2MM model. If a burst contains more than one state, it is assigned as being dynamic. (E) Dwell-based 2D E-S scatter plot. Dwells are colored on the basis of the assigned state of the chosen mpH2MM model. The dwells were corrected for leakage, direct excitation and the γ-factor. Black dots in A, D and E represent the average value of each state.
