## Supplementary File 5 for "The substrate-binding domains of the osmoregulatory ABC importer OpuA transiently interact"

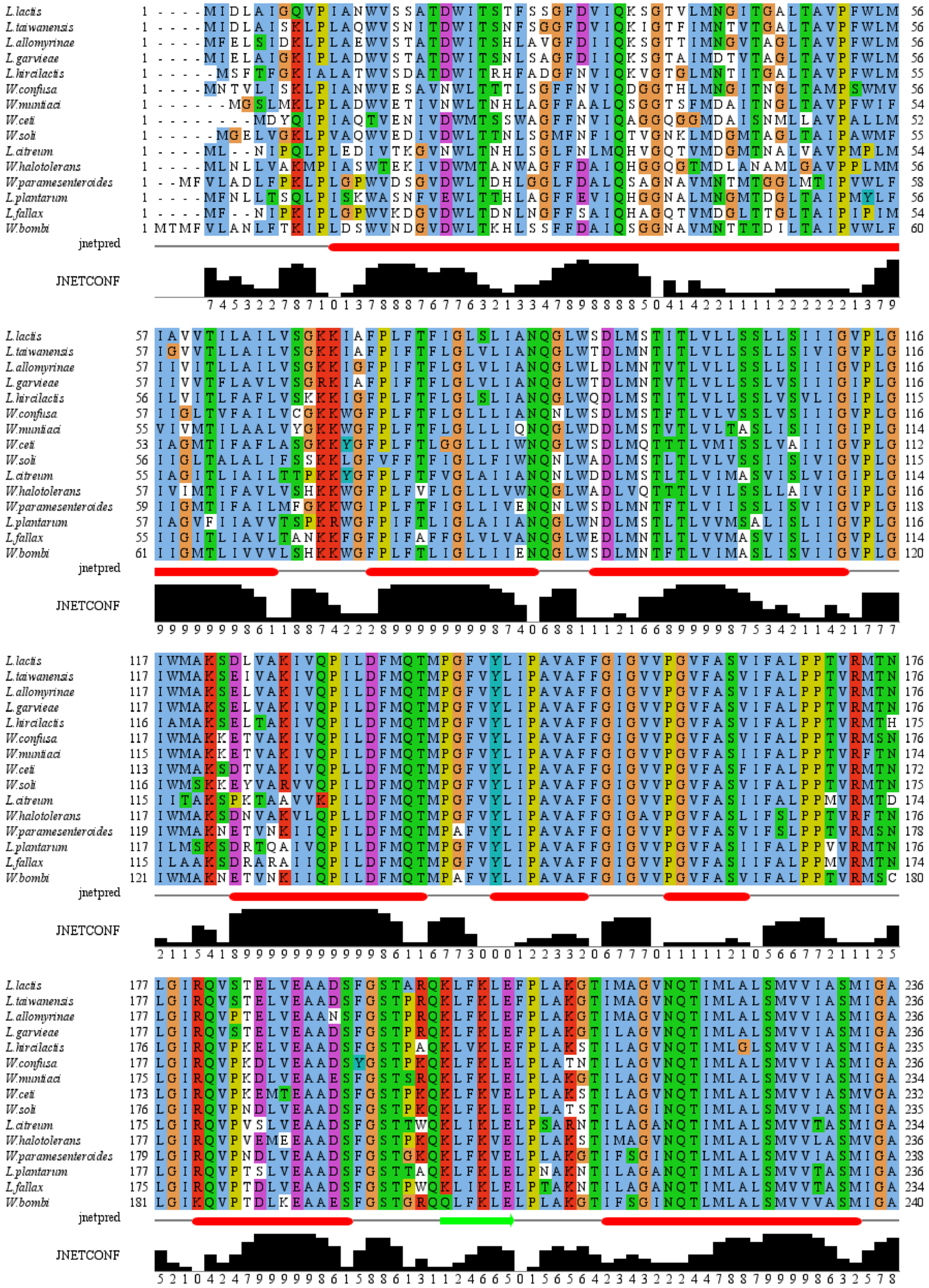

L.lactis
L.taiwanensis
L.allomyrinae
L.garvieae
L.hircilactis
W.confusa
W.muntiaci
W.ceti
W.soli
L.citreum
W.halotolerans
W.paramesenteroides
L.plantarum
L.fallax
W.bombi
jnetpred
JNETCONF

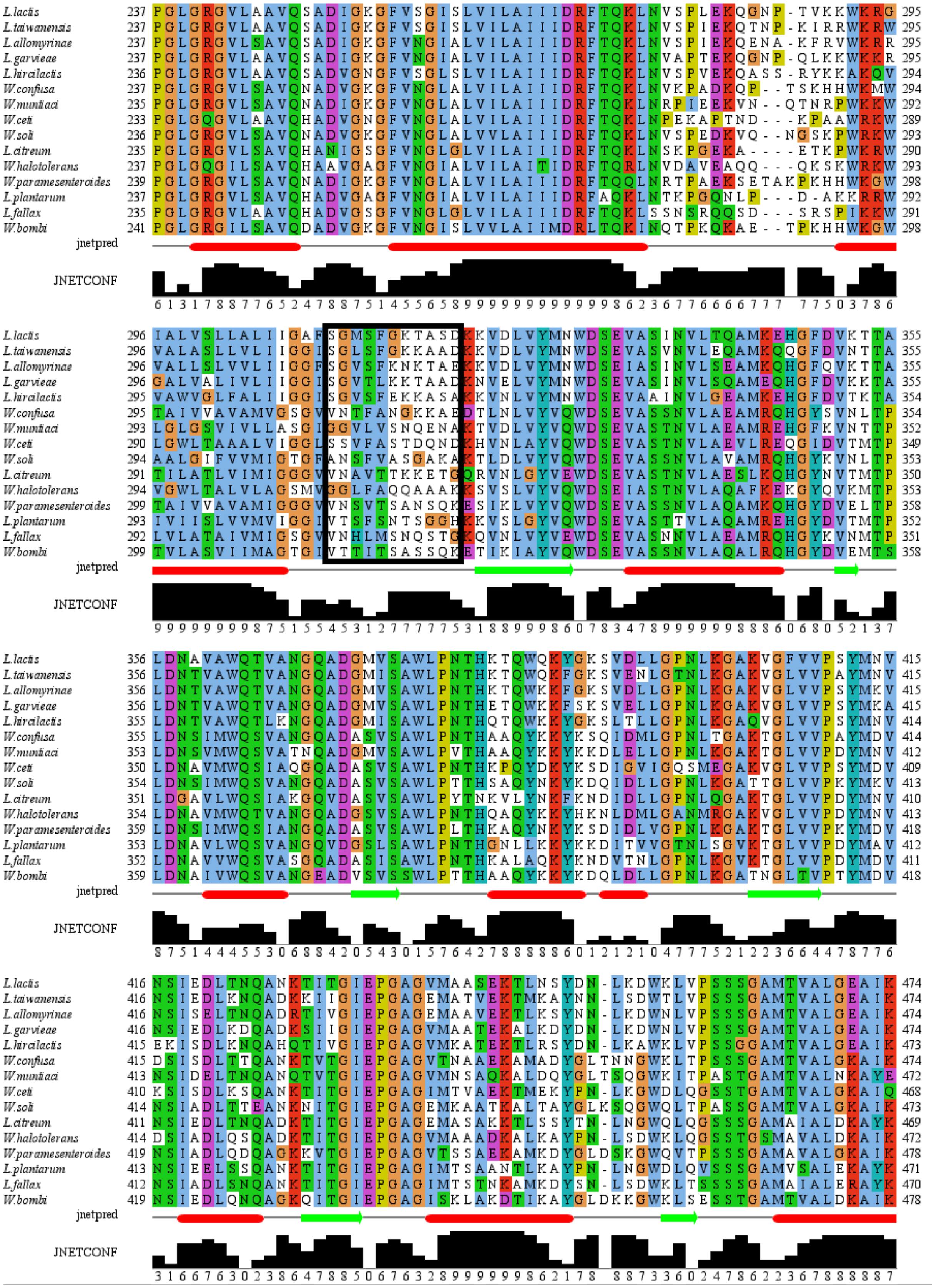
L.lactis
L.taiwanensis
L.allomyrinae
L.garvieae
L.hircilactis
W.confusa
W.muntiaci
W.ceti
W.soli
L.citreum
W.halotolerans
W.paramesenteroides
L.plantarum
L.fallax
W.bombi
jnetpred
JNETCONF

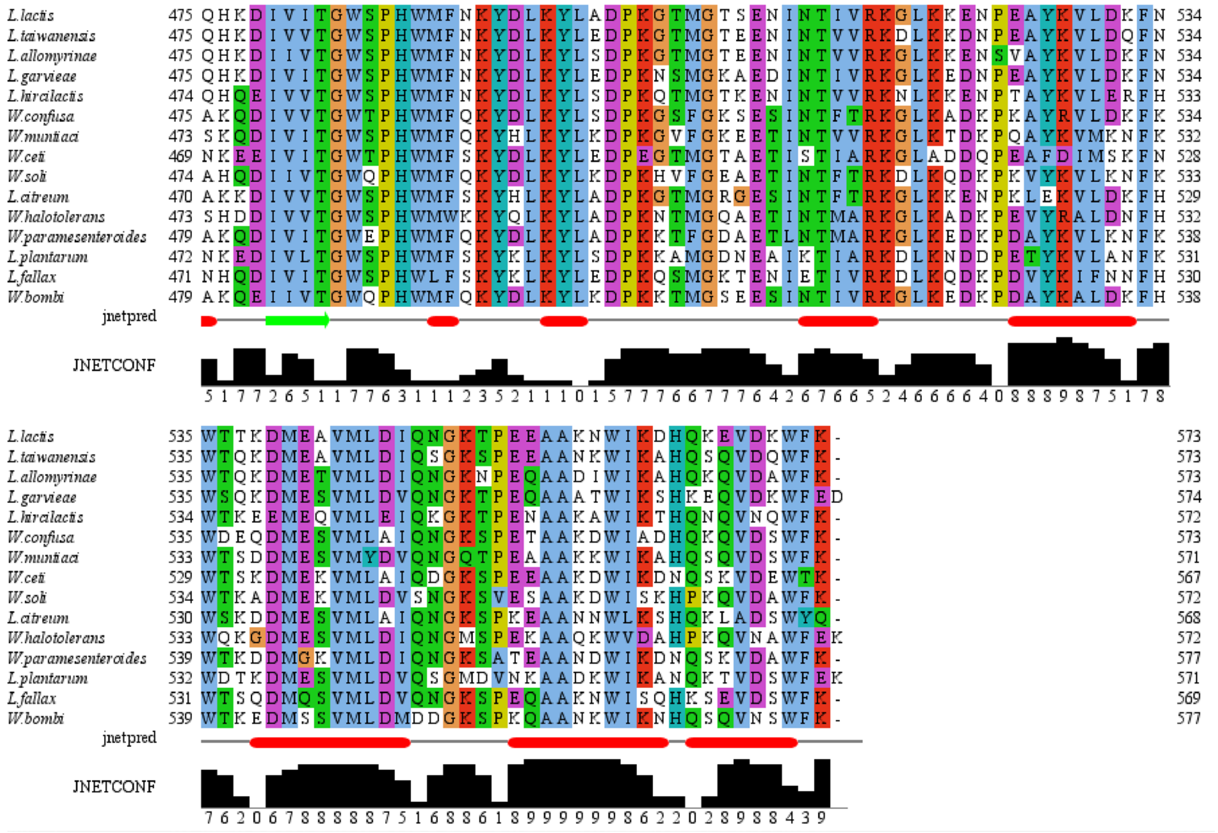

FIGURE S8. Multiple sequence alignment of close homologs of OpuA from *L. lactis*. The alignment was made by the Clustal Omega webserver using the default settings. The alignment was visualized and the JPRED secondary structure prediction were performed in Jalview (version 2.11.2.6) [3]. The black box indicates the linker region, based on OpuA from *L. lactis*. Jnetpred shows the consensus prediction of the different predictions in JPRED; α-helices are shown as red tubes and β-sheets as dark green arrows. JNETCONF is a confidence estimate of the jnetpred prediction. NCBI reference sequence IDs are: *L. lactis*, WP_003130445.1; *L. taiwanensis*, WP_205272268.1; *L. allomyrinae*, WP_120771492.1; *L. garvieae*, WP_004257235.1; *L. hircilactis*, WP_153496572.1; *W. confusa*, WP_199402959.1; *W. muntiaci*, WP_187387617.1;, *W. ceti*, WP_213409458.1; *W. soli*, WP_147152447.1; *L. citreum*, WP_040177303.1; *W. halotolerans*, WP_022790870.1; *W. paramesenteroides*, WP_150189650.1; *L. plantarum*, WP_068161132.1; *L. fallax*, WP_010007192.1; *W. bombi*, WP_092461275.1.
