## Supplementary File 6 for "The substrate-binding domains of the osmoregulatory ABC importer OpuA transiently interact"

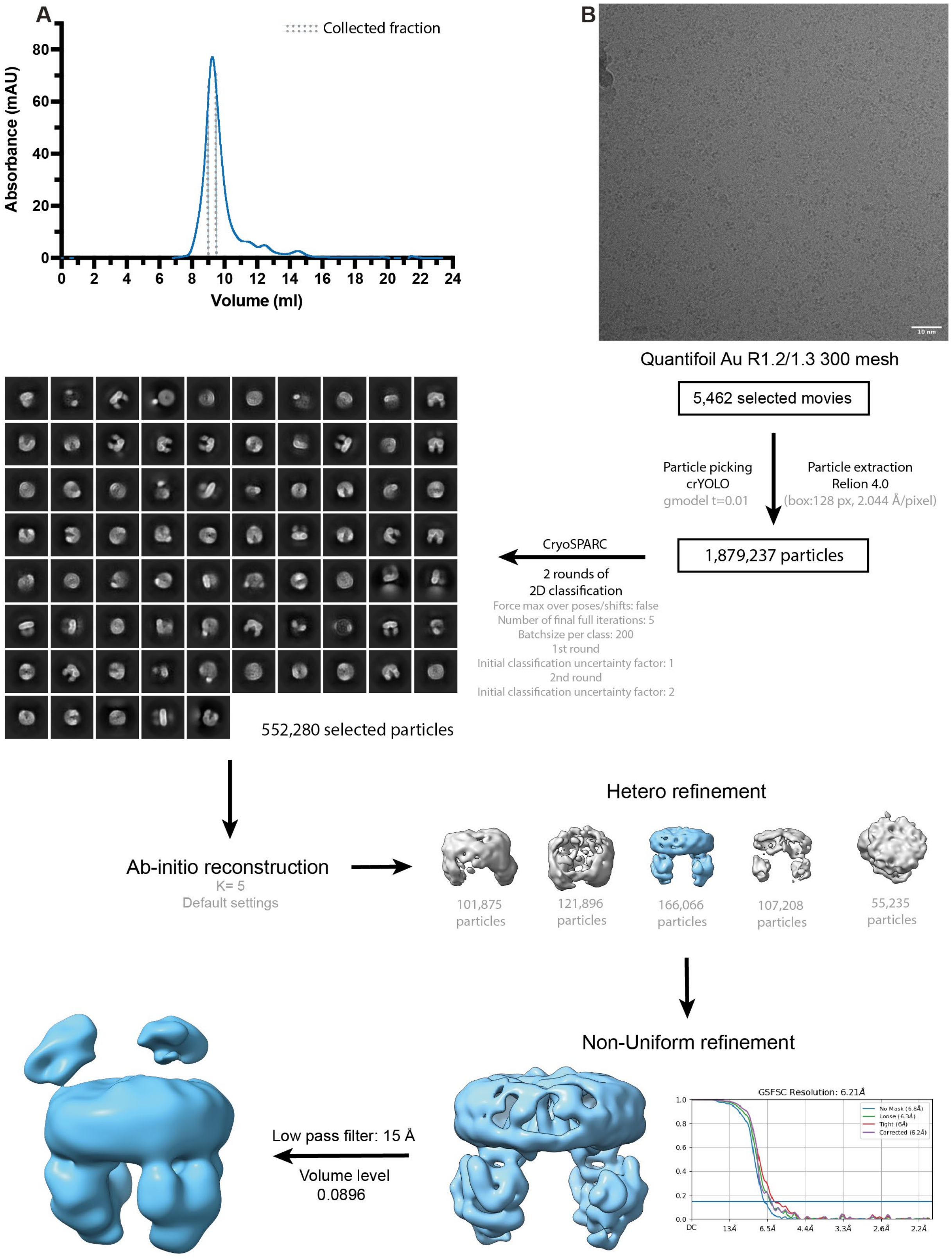

A
Collected fraction
Absorbance (mAU)
80
60
40
20
0
0 2 4 6 8 10 12 14 16 18 20 22 24
Volume (ml)
B
10 nm
Quantifoil Au R1.2/1.3 300 mesh
5,462 selected movies
Particle picking
crYOLO
gmodel t=0.01
Particle extraction
Relion 4.0
(box:128 px, 2.044 Å/pixel)
1,879,237 particles
CryoSPARC
2 rounds of
2D classification
Force max over poses/shifts: false
Number of final full iterations: 5
Batchsize per class: 200
1st round
Initial classification uncertainty factor: 1
2nd round
Initial classification uncertainty factor: 2
552,280 selected particles
Ab-initio reconstruction
K= 5
Default settings
Hetero refinement
101,875 particles
121,896 particles
166,066 particles
107,208 particles
55,235 particles
Non-Uniform refinement
GSFSC Resolution: 6.21Å
No Mask (6.8Å)
Loose (6.3Å)
Tight (6Å)
Corrected (6.2Å)
1.0
0.8
0.6
0.4
0.2
0.0
DC 13Å 6.5Å 4.4Å 3.3Å 2.6Å 2.2Å
Low pass filter: 15 Å
Volume level
0.0896

FIGURE S1. Image processing of OpuA wild-type in MSP1E3D1 nanodiscs in 20 mM HEPES-K pH 7.0, 50 mM KCl. (A) Representative elution profile from Superdex 200 increase 10/300 GL size-exclusion column. The collected fraction for cryo-EM sample preparation is indicated by dashed lines. (B) Representative micrograph at 1.3 µm defocus followed by detailed overview of the image processing leading to the map used. Abbreviations: gmodel t = crYOLO general model and K = number of classes.

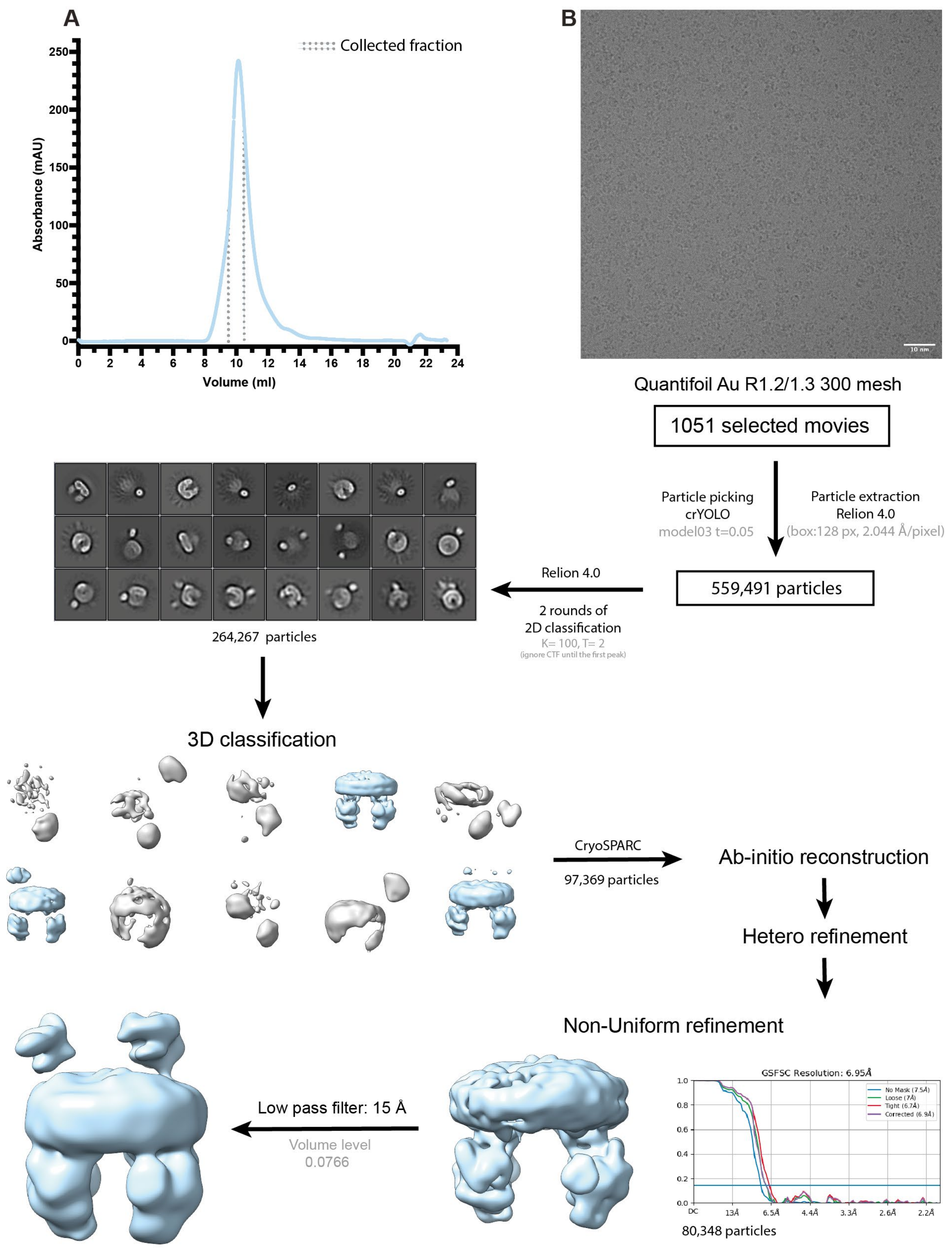
A
Collected fraction
Absorbance (mAU)
250
200
150
100
50
0
0
2
4
6
8
10
12
14
16
18
20
22
24
Volume (ml)
B
10 nm
Quantifoil Au R1.2/1.3 300 mesh
1051 selected movies
Particle picking
crYOLO
model03 t=0.05
Particle extraction
Relion 4.0
(box:128 px, 2.044 Å/pixel)
559,491 particles
Relion 4.0
2 rounds of
2D classification
K= 100, T= 2
(ignore CTF until the first peak)
264,267 particles
3D classification
CryoSPARC
97,369 particles
Ab-initio reconstruction
Hetero refinement
Non-Uniform refinement
GSFSC Resolution: 6.95Å
No Mask (7.5Å)
Loose (7Å)
Tight (6.7Å)
Corrected (6.9Å)
1.0
0.8
0.6
0.4
0.2
0.0
DC
13Å
6.5Å
4.4Å
3.3Å
2.6Å
2.2Å
80,348 particles
Low pass filter: 15 Å
Volume level
0.0766

FIGURE S2. Image processing of OpuA wild-type in MSP1E3D1 nanodiscs in 20 mM HEPES-K pH 7.0, 100 mM KCl. (A) Representative elution profile from Superdex 200 increase 10/300 GL size-exclusion column. The collected fraction for cryo-EM sample preparation is indicated by dashed lines. (B) Representative micrograph of the sample at 1.3 µm defocus, followed by detailed overview of the image processing leading to the map used. Abbreviations: model03= in-house trained crYOLO model generated by Sikkema et al. [2], K = number of classes and T= *tau_fudge.*
